# Metabolic pathway competition sensitizes thioredoxin reductase-depleted *Candida albicans* to cell wall stress and antifungals

**DOI:** 10.64898/2025.12.02.691982

**Authors:** Wanjun Qi, Udita Roy, Chunhui Cai, Maikel Acosta-Zaldívar, Jossalyn Mascio, John M. Asara, José F. Fierro, María T. Andrés, Liang Sun, Julia R. Köhler

## Abstract

*Candida albicans* is the most common invasive human fungal pathogen. Each of the currently available 3 antifungal drug classes is limited by toxicities, drug interactions or inadequate penetration of critical tissues. We discovered a connection between *C. albicans* TOR and thioredoxin reductase, Trr1. Fungal and human thioredoxin reductases diverge fundamentally and are essential for repair of oxidatively damaged macromolecules, so they may be promising drug targets. We found that Trr1 is required for *C. albicans*’ growth at human body temperatures, hyphal growth regulation and management of oxidative and cell wall stress. Its depletion sensitizes *C. albicans* to AmphotericinB and to an echinocandin. *TRR1*-depleted cells have lower cell wall glucan content. Metabolomics experiments highlighted their perturbed pentose phosphate pathway (PPP). They have sharply elevated activity of glucose-6-phosphate dehydrogenase, the first, rate-limiting enzyme of the PPP. A key glycolytic, gluconeogenic and UDP-glucose biosynthetic enzyme each show decreased activity. UDP-glucose is the substrate of cell wall glucan-producing enzymes. We propose that Trr1-depleted cells’ accelerated glucose-6-phosphate flux into the PPP, driven by demand for NADPH reducing equivalents, diminishes glucose-6-phosphate availability for UDP-glucose production and hence for cell wall construction, weakening the wall. Strikingly dysregulated signaling pathways in Trr1-depleted cells contribute to their stress hypersensitivities.

## Introduction

All organisms in aerobic environments experience oxidative stress. The mitochondrial electron transport chain leaks electrons that reduce oxygen prematurely to generate superoxide. Fatty acid oxidation in peroxisomes and protein folding in the endoplasmic reticulum generate H_2_O_2_. Pathogens like *Candida albicans* must additionally cope with exogenous superoxide released by the host immune system. Essential mitochondrial and cytoplasmic superoxide dismutases convert the superoxide anion into H_2_O_2_ and elemental oxygen _^1,2^_. Peroxidases and catalases reduce H_2_O_2_ to elemental oxygen and water _^3^_. H_2_O_2_ diffuses freely through membranes and if it reacts with Fe_^2+^_, a component of many enzymes’ active sites, the extremely reactive hydroxyl radical is generated which oxidizes surrounding macromolecules including proteins, nucleic acids and membranes.

Oxidative damage to macromolecules is repaired by specialized thiol-disulfide redox systems: the abundant tripeptide glutathione, the small protein thioredoxin as well as the peroxidases, the most important of which are the peroxiredoxins ^4-8^. The oxidized reactants in these repair systems are reduced and thereby reactivated by glutathione- and thioredoxin reductases using NADPH as the ultimate electron donor. Peroxiredoxins are also recycled via the thioredoxin-thioredoxin reductase system ^9^.

Thioredoxin reductases are protein disulfide oxido-reductases ^10^ that recycle oxidized thioredoxin to its active reduced state. Thioredoxin reductase of humans and other higher eukaryotes is 20 kD larger and has a different structure than the thioredoxin reductases of bacteria, plants, fungi, and protozoal parasites ^11^. Because of their critical role in cellular redox homeostasis and their divergence from the human enzyme, thioredoxin reductases of fungal and bacterial pathogens have been proposed as promising drug targets ^12-16^.

In a previous study, we found that *TRR1*, the gene encoding *C. albicans*’ thioredoxin reductase, is upregulated 2.6-fold higher during oxidative stress in wild type (WT) than in *TOR1*^*Del381*^ mutants in the Target of Rapamycin 1 kinase ^17^. Others previously found *TRR1* was required in *C. albicans* pathogenesis because its heterozygous deletion mutants were significantly less virulent in a systemic mouse infection model than their wild type (WT) progenitor ^18^. On this basis, we further investigated its role.

## Results

### The gene encoding thioredoxin reductase Trr1 is conserved among fungi and divergent from that of humans

We found that thioredoxin reductases from several human-pathogenic fungal species, including *C. albicans*, share highly conserved predicted primary amino acid sequences. The human homolog, as expected, shows a very different protein structure (SI Appendix, Fig. S1). To investigate essentiality of Trr1 in *C. albicans*’ growth and stress responses, we engineered *C. albicans* strains in which one *TRR1* allele is deleted while the remaining allele is expressed from repressible *tetO*.

In addition to the thioredoxin system, *C. albicans* uses the tripeptide glutathione to reduce proteins damaged by oxidation ^19^. The first step in glutathione biosynthesis is performed by *γ*-glutamyl cysteine synthase, encoded by *GCS1* ^19,20^. Deletion of *GCS1* results in auxotrophy for glutathione; *gcs1* null mutants grown without glutathione are avirulent in an intravenous mouse model of infection ^20^. Several human-pathogenic unicellular parasites like *Entamoeba histolytica* lack glutathione ^21^. To rigorously compare the importance of the thioredoxin- and glutathione antioxidant systems for *C. albicans’* stress management, we constructed *gcs1* mutants with the same genetic perturbation as in *TRR1*, deletion of one allele and control of the other by repressible *tetO*. For further phenotypic analysis, we also constructed *gcs1/gcs1* null mutants and *gcs1/gcs1 trr1/tetO-TRR1* mutants in the same genetic background.

### *TRR1* is essential for *C. albicans*’ growth at host temperatures

Depleting *TRR1* from *tetO* resulted in a substantial growth defect of *C. albicans* on solid media under optimal in vitro conditions including *C. albicans’* preferred temperature, 30^°^ C. (Fig. 1A and SI Appendix, Fig. S2A). Correspondingly, TOR Complex 1 pro-anabolic activity was decreased in *TRR1*-depleted cells, reflected in the phosphorylation state of ribosomal protein S6 in vehicle (pS6) (Fig. 1B). *TRR1-*depleted cells’ growth defect was severe at 37^°^. At 40^°^, a temperature actively produced during infection by human innate immune responses, these cells failed to grow (Fig. 1A and SI Appendix, Fig. S2A). Depletion of *GCS1* from *tetO* also sensitized cells to temperatures above 30^°^, though their growth defect was substantially less than that of *TRR1*-depleted cells (Fig. 1A and SI Appendix, Fig. S2A). Hence Trr1 is essential for *C. albicans* growth at host temperatures, while Gcs1 contributes to temperature tolerance.

**Figure 1.**
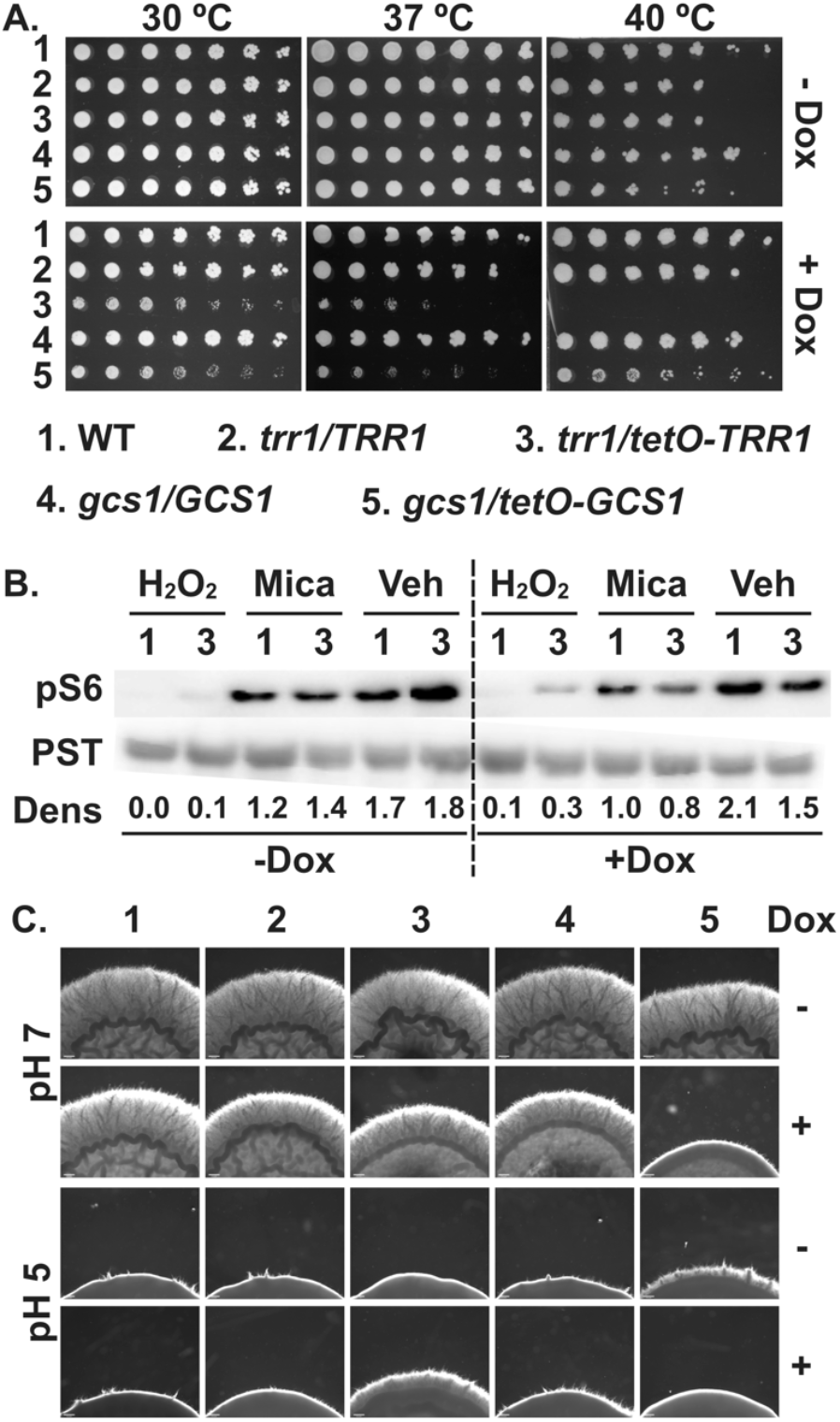
*TRR1* is essential at host body temperature and required for normal TOR signaling and filamentous growth regulation. **A**. Cell dilutions (left to right) of indicated genotypes spotted onto synthetic complete (SC) plates without or with 1 μg/ml doxycycline (Dox) and grown at 30^°^, 37^°^ and 40^°^ C for 2 days. **B**. Cells of indicated genotypes pre-grown in YPD without or with 300 ng/ml doxycycline for 3.5 h, then inoculated into YPD containing 2.5 mM H_2_O_2_, 10 ng/ml micafungin (Mica) or vehicle (Veh, H_2_O), without or with 300 ng/ml doxycycline. Cells harvested at 5 min for H_2_O_2_ and 30 min for micafungin and vehicle. Extract blotted and probed against phosphorylated S6 (pS6) and PSTAIRE (PST) as loading control. Densitometry (Dens): intensity ratio between pS6 and PST. **C**. Cell suspensions of indicated genotypes were spotted at equidistant points around the perimeter of RPMI plates, without or with 1 μg/ml doxycycline, and grown at 37^°^ for 2 days. Size bar 200 μm. Strains are 1. WT (JKC915); 2. *trr1/TRR1* (JKC2989); 3. *trr1/tetO-TRR1* (JKC2997); 4. *gcs1/GCS1* (JKC3001); 5. *gcs1/tetO-GCS1* (JKC3009).

### Hyphal morphogenesis is dysregulated by *TRR1-* and stunted by *GCS1* depletion

Frequent morphogenetic switches, i.e. production of hyphal or pseudohyphal filaments from yeast in response to external conditions as well as constitutive production of lateral yeast on filaments, contribute to virulence of *C. albicans* ^22-25^. Previously, changes in glutathione abundance and oxidation state were reported during *C. albicans* filamentation ^26,27^. We found that Trr1 was required for appropriate regulation of morphogenesis: at hyphae-inducing pH 7, *TRR1* depletion resulted in slightly decreased filamentation while at hyphae-repressing pH 5, filamentation was inappropriately increased (Fig. 1C). *GCS1* depleted cells formed substantially less hyphae than WT at neutral pH (Fig. 1C). Together with temperature phenotypes of *TRR1-*depleted cells, their dysregulated filamentation suggests important roles for Trr1 in *C. albicans’* virulence for a human host.

### Trr1 is essential for *C. albicans*’ oxidative stress management

Human phagocytes release superoxide anion onto intracellular *C. albicans* cells or extracellularly in Neutrophil Extracellular Traps. Superoxide is dismutated into H_2_O_2_ and O_2_ by fungal superoxide dismutases ^1,7^, and peroxide stress management requires thioredoxin ^5,28^. Since thioredoxin reductase recycles oxidized thioredoxin, we tested the role of Trr1 in oxidative stress. *TRR1*-depleted cells were severely hypersensitive to superoxide stress induced by plumbagin exposure and to peroxide stress induced by H_2_O_2_ (Fig. 2A and SI Appendix, Fig. S2B). Lack of one *TRR1* allele was sufficient to cause a growth defect during peroxide but not superoxide exposure. *GCS1* loss-of-function phenotypes during exposure to these oxidative stressors were overall weaker, with no apparent haploinsufficiency (Fig. 2A and SI Appendix, Fig. S2B).

**Figure 2.**
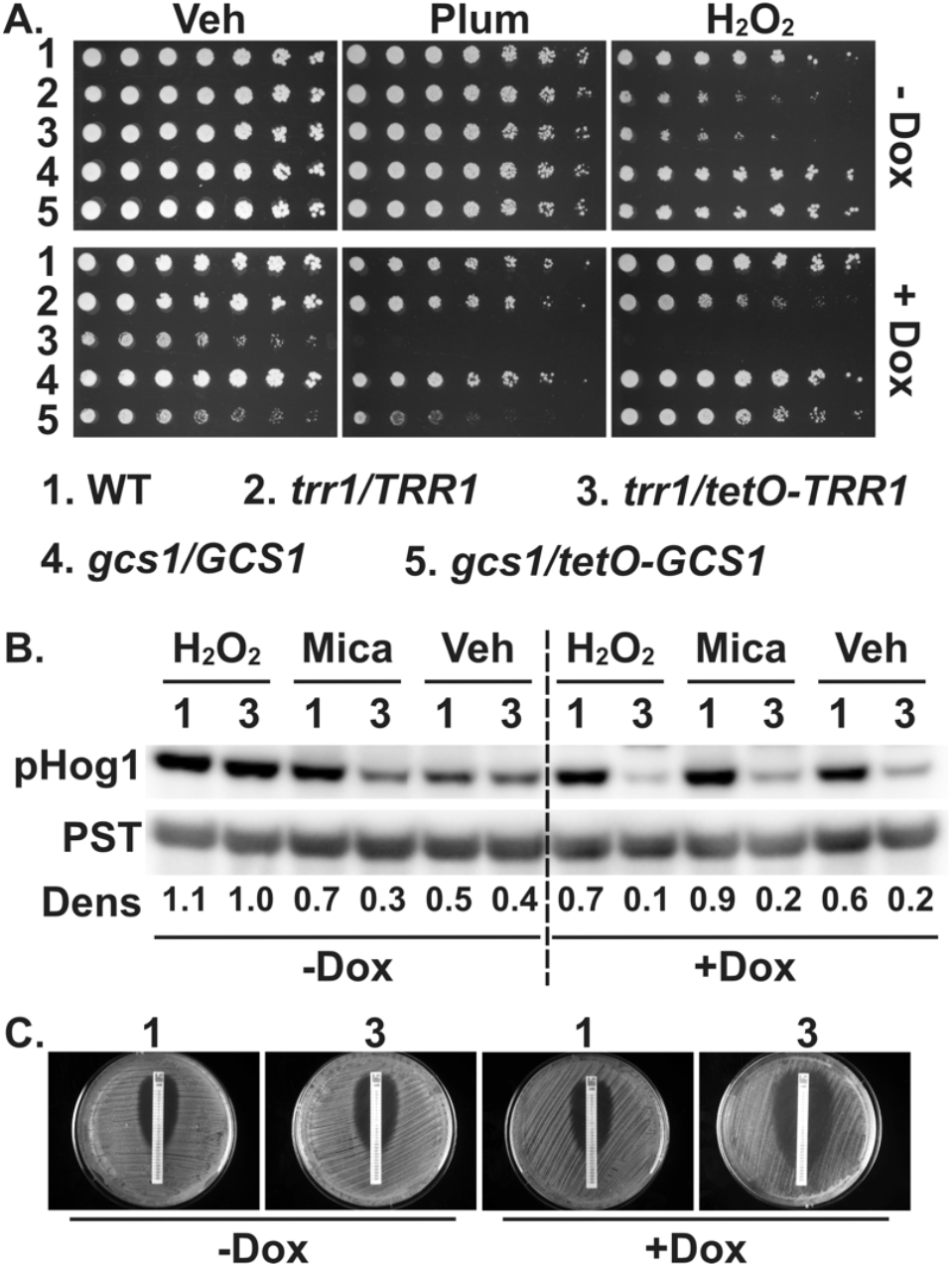
*TRR1* is required for oxidative stress management. **A**. Cell dilutions (left to right) of indicated genotypes spotted onto SC plates without or with 1 μg/ml doxycycline (Dox) containing vehicle (Veh, H_2_O), 15 μM plumbagin (Plum) or 2.5 mM H_2_O_2_, grown at 30_^°^_ C for 2 days. **B**. Cells of indicated genotypes grown as in Fig. 1B. Extract blotted and probed against phosphorylated Hog1 (pHog1) and PSTAIRE (PST) as loading control. Densitometry (Dens): intensity atio between pHog1 and PST. **C**. AmphotericinB Etest. Cells of indicated genotypes were swabbed on RPMI pH 7 plates without or with 1 μg/ml doxycycline (Dox) and incubated at 35° for 24 h with Amphotericin B (AMB 0.002-32 mg/L) Etest strips. Strains are 1. WT (JKC915); 2. *trr1/TRR1* (JKC2989); 3. *trr1/tetO-TRR1* (JKC2997);4. *gcs1/GCS1* (JKC3001); 5. *gcs1/tetO-GCS1* (JKC3009).

We examined the response of the oxidative Stress-Activated Protein Kinase Hog1 ^29,30^ to loss of Trr1 activity. *TRR1-*depleted cells had consistently diminished HOG pathway activation as reflected in the phosphorylation state of Hog1, without or with external stressors (Fig. 2B).

The broad-spectrum, candidacidal antifungal agent AmphotericinB induces oxidative stress in *C. albicans* ^31,32^. We questioned whether loss of Trr1 activity sensitizes *C. albicans* cells to AmphotericinB. *TRR1-*depleted cells had a lower AmphotericinB minimal inhibitory concentration (0.008 μg/ml) than WT (0.047 μg/ml) in the Etest assay (Fig. 2C), highlighting a drug target potential of Trr1 for use in combination with lower and hence less toxic AmphotericinB doses.

### *TRR1* depletion potentiates cell wall stress

The concepts of stress cross-protection and of anticipatory stress responses predict that exposure to one stress primes cells to resist another stress ^33,34^. But pre-existing oxidative stress in cells depleted for *TRR1* did not protect but instead sensitized *C. albicans* to all 3 tested cell wall stressors: micafungin which inhibits biosynthesis of a beta-1,3-glucan, as well as calcofluor white and Congo Red which bind to and disrupt chitin fibrils (Fig. 3A and SI Appendix, Fig. S2C).

**Figure 3.**
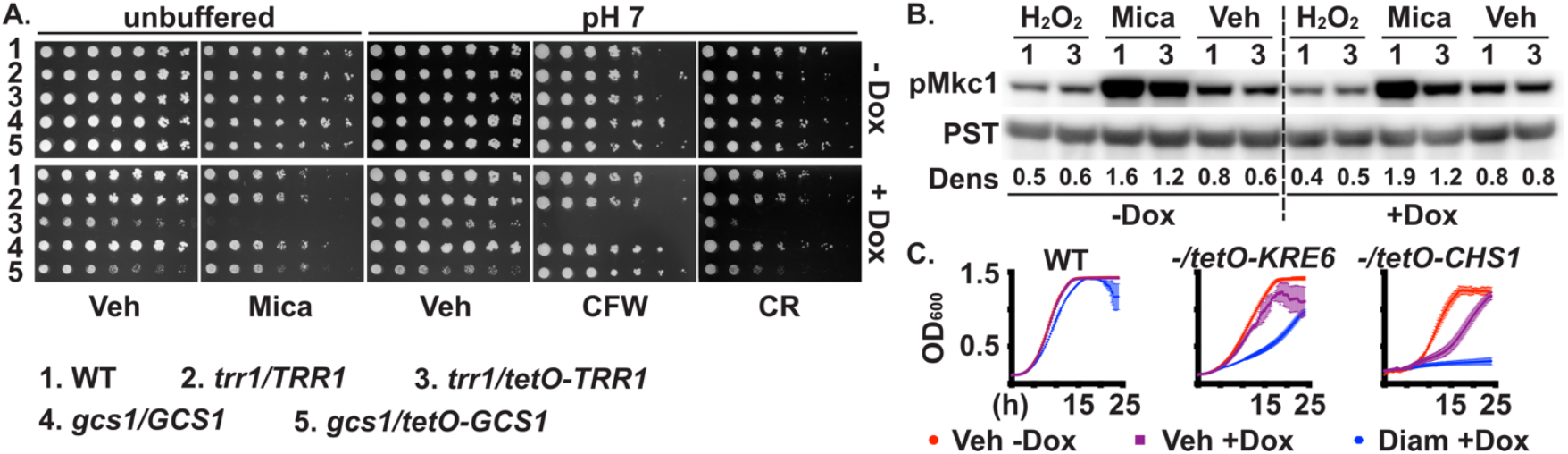
*TRR1* is required for cell wall stress management. **A**. Cell dilutions (left to right) of indicated genotypes spotted onto SC plates containing vehicle (Veh, H_2_O) or 10 ng/ml micafungin (Mica), and SC pH7 plates containing vehicle (Veh, H_2_O), 10 μg/ml calcofluor white (CFW) or 10 μg/ml Congo Red (CR), without or with 1 μg/ml doxycycline (Dox), grown at 30^°^ for 2 days. **B**. Cells of indicated genotypes grown as in Fig. 1B. Extract blotted and probed against phosphorylated Mkc1 (pMkc1) and PSTAIRE (PST) as loading control. Densitometry (Dens): intensity ratio between pMkc1 and PST. **C**. Growth of indicated genotypes in SC media with vehicle (Veh, DMSO) or 0.5 mM diamide (Diam), without or with 500 ng/ml doxycycline (Dox). Strains are 1. WT (JKC915); 2. *trr1/TRR1* (JKC2989); 3. *trr1/tetO-TRR1* (JKC2997); 4. *gcs1/GCS1* (JKC3001); 5. *gcs1/tetO-GCS1* (JKC3009). *-/tetO-KRE6* (*kre6/tetO-KRE6*, JKC2198); *-/tetO-CHS1* (*chs1/tetO-CHS1*, JKC2212).

Cells lacking Trr1 were notably defective in cell wall stress signaling. In response to micafungin, *TRR1*-depleted cells failed to appropriately activate the cell wall integrity (CWI) signaling pathway, as indicated by the Mkc1 phosphorylation state ^35-37^ (Fig. 3B). Of note, when *tetO*-controlled *TRR1* was induced in the absence of doxycycline, CWI pathway activation was also weaker compared to the WT (Fig. 3B), suggesting physiologically controlled *TRR1* expression is required for appropriate CWI signaling.

### Cell wall- and oxidative stress potentiate each other in other genetic backgrounds

Since genetic loss of Trr1 potentiated cell wall stress, we asked whether this effect might be generalizable to combinations of cell wall- and thiol redox stress. To genetically induce cell wall stress, we used our *C. albicans* mutants in which the only allele of genes encoding the essential chitin synthase Chs1 or the beta-1,6-glucan synthase Kre6 is expressed from repressible *tetO* ^38^. During *CHS1* or *KRE6* repression, these cells were hypersensitive to the thiol-disulfide oxidative stress inducer diamide ^39^ (Fig. 3C). Mutual potentiation of cell wall- and thiol-oxidative stress therefore occurred in other constellations of genetic and small molecule inhibition; it was not limited to loss of *TRR1*.

To eliminate a possible effect of *tetO*-repressive doxycycline on stress phenotypes, we tested cell wall- and oxidative stress combinations against WT, *trr1/TRR1, gcs1/GCS1* and *trr1/TRR1 gcs1/GCS1* hemizygous mutants, relying on haploinsufficiency to test loss-of-function effects.

Concentrations of micafungin and H_2_O_2_ that alone had only mild effects on all genotypes tested, had inhibitory effects on hemizygous strains in combination (SI Appendix, Fig. S2D). Loss of one *TRR1* allele had a stronger effect than loss of one *GCS1* allele, while the *trr1/TRR1 gcs1/GCS1* double hemizygote had moderately increased sensitivity (SI Appendix, Fig. S2D). Taken together, these findings of mutual potentiation of oxidative- and cell wall stress suggest that low-toxicity drug combinations targeting Trr1 and cell wall biosynthesis might achieve high potency against *C. albicans*.

### Growth on lactate exacerbates *TRR1*-depleted cells’ hypersensitivity to cell wall stress

The inner layer of the fungal cell wall consists of an elastic mesh of interwoven polysaccharides. This mesh provides the mechanical strength to contain the turgor pressure exerted by the cytoplasm on the plasma membrane, and withstand mechanical stress from outside the cell ^40^. Beta-1,3- and beta-1,6-glucan together comprise 95% of the rigid components of the *C. albicans* cell wall and 64% of its mobile components according to a recent NMR analysis, while chitin contributes 5% to the rigid molecules ^41^.

A pivotal metabolic intermediate in cell wall construction and oxidative stress management is glucose-6-phosphate (glucose-6-P). As the substrate of glucose-6-phosphate dehydrogenase (G-6-PD), the first, rate-limiting enzyme of the pentose phosphate pathway (PPP), it provides reducing equivalents as NADPH ^42^. Transcription of *ZWF1*, encoding G-6-PD, is upregulated during oxidative stress in *C. albicans* ^43^. As the substrate of phosphoglucomutase, glucose-6-P is a precursor to UDP-glucose, the substrate of beta-1,3- and beta-1,6-glucan synthases ^38,44,45^. As the substrate of phosphohexoseisomerase, it is converted to fructose-6-phosphate to enter glycolysis.

Ene and colleagues previously found that the inner cell wall layer of *C. albicans* cells grown on lactate is significantly thinner than that of cells grown on glucose ^46,47^. We considered whether a dearth of glucose-6-P during lactate growth might slow UDP-glucose production such that beta-1,3- and beta-1,6-glucan synthases lack their substrate. To test this idea, we grew WT and *trr1* mutant cells on lactate, requiring them to produce glucose-6-P by gluconeogenesis. At a low micafungin concentration of 10 ng/ml, which did not inhibit WT cells’ growth on lactate, *TRR1*-depleted cells had a more severe growth defect on lactate than on glucose (Fig. 4A). Hence, metabolic pathway competition for glucose-6-P might contribute to *TRR1* cells’ hypersensitivity to cell wall stress.

**Figure 4.**
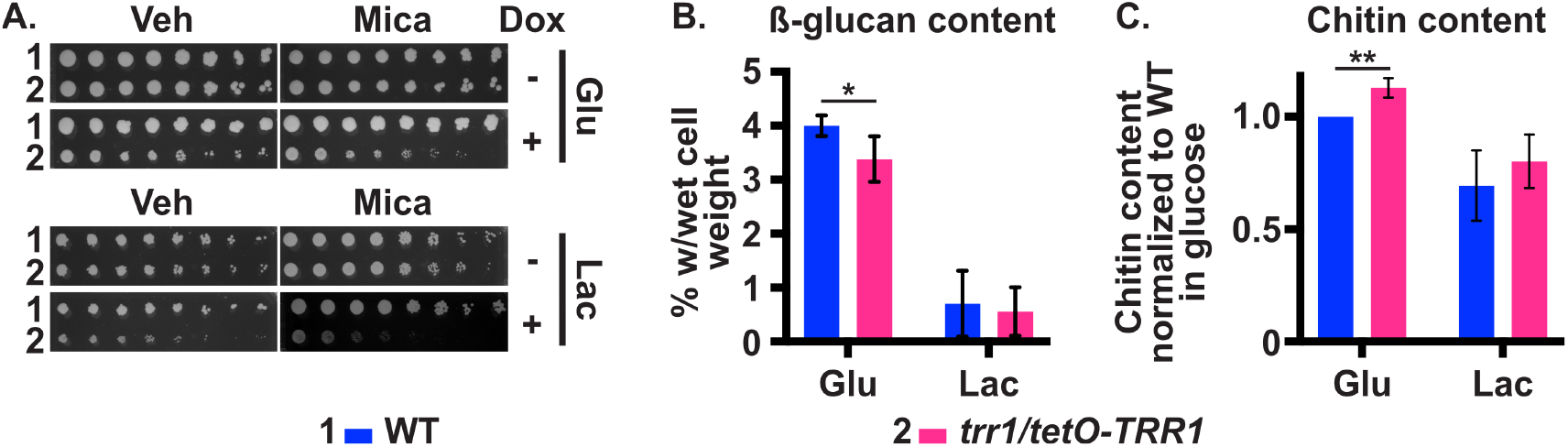
*TRR1*-depleted cells’ hypersensitivity to cell wall stress is exacerbated on a non-glucose carbon source while they show a decreased cell wall glucan content. **A**. Cell dilutions (left to right) of indicated genotypes spotted onto SC plates containing glucose (Glu) or lactate (Lac), vehicle (Veh, H_2_O) or 10 ng/ml micafungin (Mica), without or with 300 ng/ml doxycycline (Dox), grown at 30^°^ for 2 days. **B**. ß-glucan content of wild type (WT) and *trr1/tetO-TRR1* cells grown on SC plates containing 1 μg/ml doxycycline, with either 2% glucose or 2% lactate. **C**. Chitin content of cells in panel B. Strains are 1. WT (JKC915); 2. *trr1/tetO-TRR1* (JKC2997).

### *TRR1* depletion leads to decreased glucan cell wall content

We examined whether *TRR1* depletion affected cell wall glucan content: might *TRR1*-depleted cells prioritize flux of glucose-6-P into the PPP over its use in cell wall construction? In fact, glucan content of glucose-grown *TRR1*-depleted cells’ walls was significantly lower than that of WT cells (Fig. 4B). Cell wall glucan content of lactate-grown cells of both genotypes was dramatically lower than that of the same cells grown on glucose, with *TRR1*-depleted lactate-grown cells showing a statistically non-significant trend of lower glucan content than WT (Fig. 4B).

Beta-1,3-glucan synthesis inhibition is known to induce compensatory increased chitin biosynthesis ^48,49^. The same cultures whose cell wall glucan content was measured, were assayed for chitin content as in ^38^. During growth in glucose, chitin content of *TRR1*-depleted cells’ walls was significantly higher than that of WT (Fig. 4C). The increase of stabilizing chitin suggests a cell wall stress response when cells lack Trr1. This response seems partially Mkc1-independent (Fig. 3B).

### Metabolomics analysis highlights PPP perturbations in *TRR1*-depleted cells

In *S. cerevisiae* and *Caenorhabditis elegans*, oxidative stress redirects glucose-6-P flux into the PPP within seconds, increasing NADPH production ^50,51^. To test the idea that metabolic pathway competition for glucose-6-P leads to mutual potentiation of oxidative- and cell wall stress, we performed metabolomics experiments. Six independent biological replicates were obtained from WT and partially *TRR1*-depleted cells grown in 300 ng/ml doxycycline in glucose or lactate.

Hydrophilic interaction liquid chromatography-mass spectrometry (LC-MS/MS) quantitated 298 known metabolites as in ^52^; analysis of signals from polar metabolites was performed using MetaboAnalyst {Pang, 2024 #103}. Principal component analysis showed sharp separation of metabolites derived from WT- and *TRR1*-depleted cells grown in glucose; lactate-grown metabolites of the 2 strains showed closer clustering (SI Appendix, Fig. S3A). Further analysis focused mainly on glucose-grown cells.

Initial inspection of the Top 25 metabolites differing most significantly between WT and *TRR1*-depleted cells (SI Appendix, Fig. S3B) highlighted fructose-6-phosphate at the top. Fructose-6-phosphate is the 3rd metabolite in glycolysis directly below a tripartite branchpoint at glucose-6-P, which can enter 3 pathways: the PPP by G-6-PD, glycolysis by phosphohexoseisomerase, or polysaccharide and trehalose production via UDP-glucose by phosphoglucomutase (Fig. 5A).

**Figure 5.**
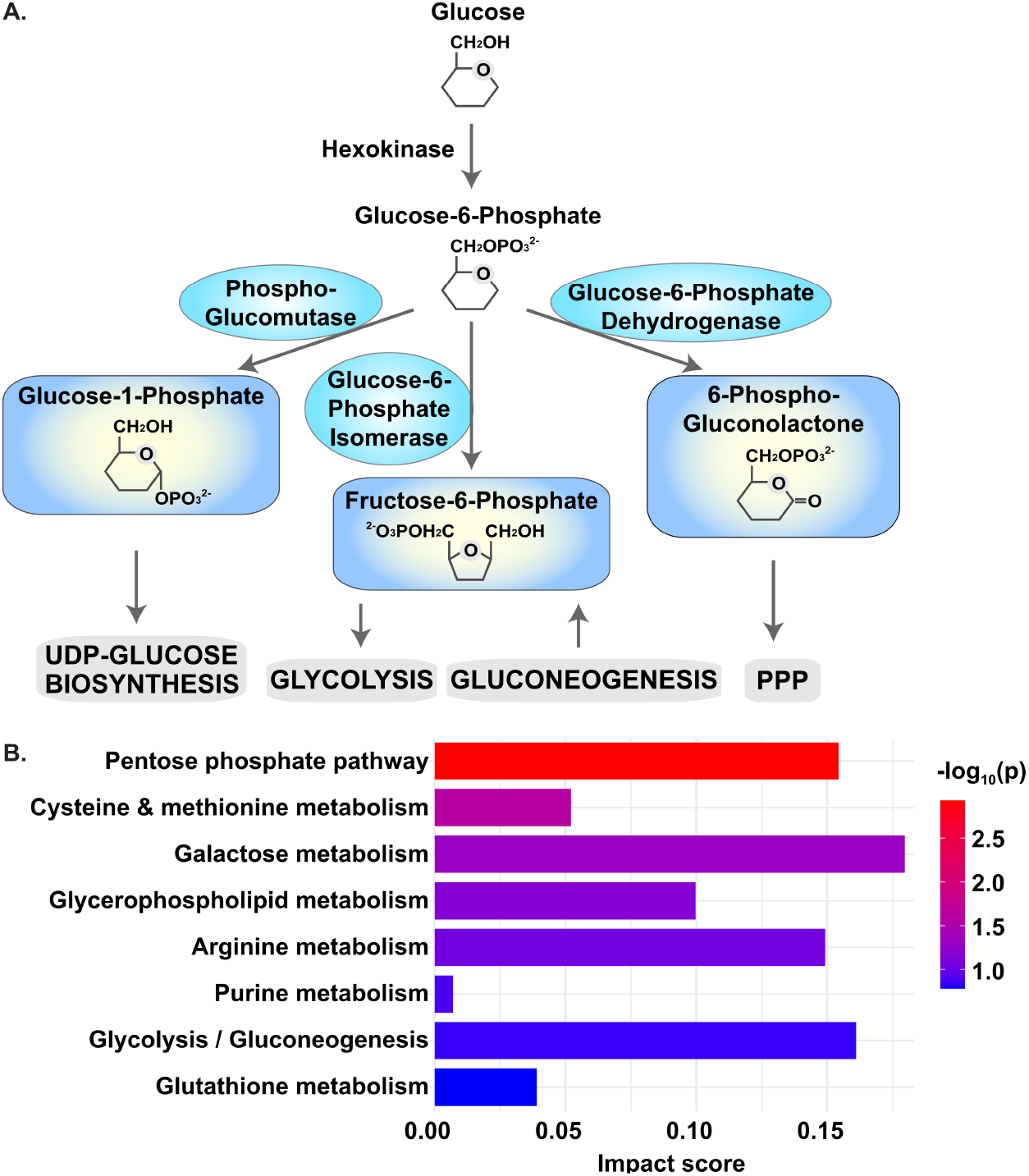
Metabolic pathway competition may decrease cell wall biosynthetic precursors in *TRR1*-depleted cells. **A**. Metabolic pathways competing for glucose-6-phosphate (PPP, pentose phosphate pathway). **B**. Pathway enrichment analysis of metabolomic perturbations of *trr1/tetO-TRR1* versus wild type cells grown in glucose with 300 ng/ml doxycycline. Analysis was performed using the MetaboAnalyst 6.0 ‘Pathway Analysis’ module based on the KEGG database, displaying significantly enriched pathways with p < 0.05 and pathway impact factor > 0.

Decreased levels of fructose-6-phosphate in *TRR1*-depleted cells suggested that glycolysis was not favored at the glucose-6-P branchpoint. Glucose-6-P itself was decreased in *TRR1*-depleted cells (SI Appendix, Fig. S3B).

We examined the differentially affected metabolic pathways in *TRR1*-depleted versus WT cells. In glucose-grown cells, the most significantly affected pathway was the PPP (Fig. 5B). Also affected was biosynthesis of sulfur-containing amino acids methionine and cysteine. We interpreted the latter as confirmatory controls of the metabolomics experiments: *C. albicans* thioredoxin, the substrate of Trr1, is known to be required for methionine biosynthesis ^5,53^. We found *TRR1*-depleted cells to be auxotrophic for methionine and cysteine (SI Appendix, Fig. S4A).

Metabolomics analysis of glucose-grown cells showed significantly lower NADPH levels in *TRR1*-depleted cells than in WT (SI Appendix, Fig. S4B). PPP intermediates like sedoheptulose-1-7-phosphate, ribose-5-phosphate and erythrose-4-phosphate were also decreased in glucose-grown *TRR1*-depleted cells (SI Appendix, Fig. S4B), raising the question whether in these cells, flux through the PPP was increased (as predicted by general regulatory responses to oxidative stress) or decreased (if diminished *TRR1* transcription and Trr1 protein levels reduced NADPH consumption).

### Glucose-6-P dehydrogenase activity is increased in *TRR1*-depleted cells

As our steady-state metabolomics experiments do not distinguish increased consumption of NADPH from its decreased production, we assayed the activity of the first and rate-limiting enzyme of the PPP, G-6-PD. We used growth conditions exactly corresponding to the metabolomics experiments. G-6-PD activity was significantly higher in *TRR1*-depleted cells than in WT in both glucose and lactate (Fig. 6A), indicating that flux through the upper PPP was increased in these cells. G-6-PD activity was lower in lactate-grown WT and *TRR1*-depleted cells, possibly reflecting these cells’ reliance on gluconeogenesis alone for their glucose-6-P supply. The inability of *TRR1*-depleted cells to appropriately manage thiol-redox stress hence mimics other oxidative stressors like peroxide exposure, in that it induces increased flux through the PPP. This is consistent with G-6-PD activation through a decreased NADPH/NADP ratio found in numerous cell types ^54^.

**Figure 6.**
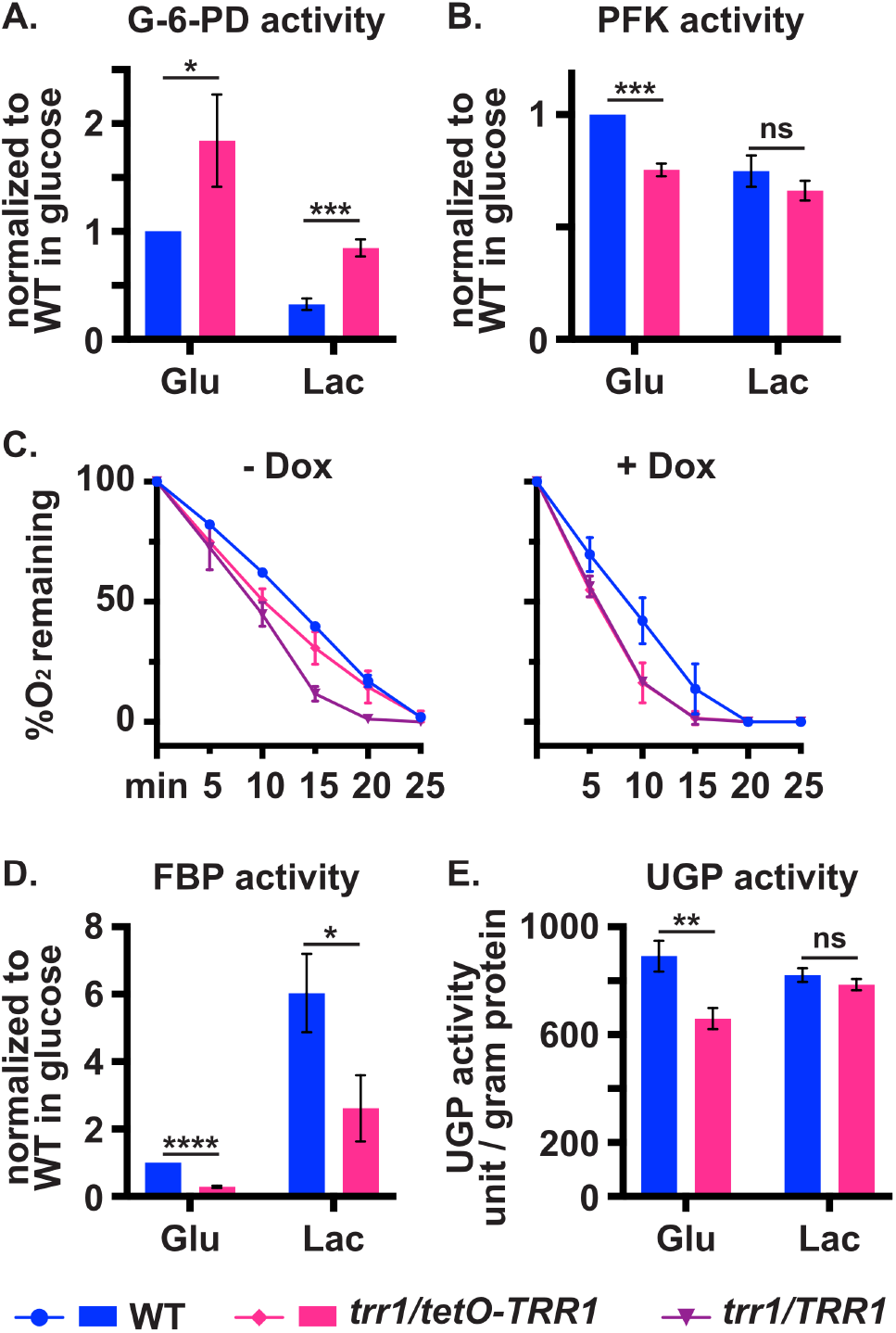
*TRR1*-depleted cells have increased PPP activity and increased oxygen consumption, but decreased glycolysis, gluconeogenesis and UDP-glucose production. Wild type and *trr1/tetO-TRR1* cells grown in glucose or lactate with 300 ng/ml doxycycline assayed for enzymatic activities of glucose-6-phosphate dehydrogenase (G-6-PD) (pentose phosphate pathway) **(A)**, phosphofructokinase (PFK) (glycolysis) **(B)**, fructose 1,6-Bisphosphatase (FBP) (gluconeogenesis) **(D)** and uridine diphosphate glucose pyrophosphorylase (UGP) (cell wall biosynthesis) **(E). C**. Oxygen (O_2_) consumption of indicated genotypes grown in glucose without or with 300 ng/ml doxycycline (Dox). Strains are WT (JKC915); *trr1/TRR1* (JKC2989); *trr1/tetO-TRR1* (JKC2997).

### Glycolysis slows while respiration accelerates in *TRR1-*depleted cells

In *S. cerevisiae*, oxidative stress increases flux of glucose-6-P into the PPP by redox inactivation of key glycolytic enzymes ^50,51,55^. Phosphofructokinase represents a key regulatory enzyme in upper glycolysis. Phosphofructokinase activity of glucose-grown *TRR1*-depleted cells was lower than that of WT, reflecting an adaptive decrease of glycolytic flux in the context of increased glucose-6-P consumption by the PPP (Fig. 6B). In lactate, both WT and *TRR1*-depleted cells had significantly reduced phosphofructokinase activity, consistent with these cells’ glucose starvation (Fig. 6B).

In aerobic environments, respiration generates ∼15 times more ATP per molecule of glucose than glycolysis. The product of glycolysis, pyruvate, enters the tricarboxylic acid cycle after conversion into acetyl-CoA, while the reduced products of the tricarboxylic acid cycle drive the electron transport chain for final reduction of elemental oxygen to water. To test whether decelerated glycolysis in *TRR1*-depleted cells ultimately leads to decreased respiration, we measured oxygen consumption of WT, *trr1/TRR1* hemizygous, and *trr1/tetO-TRR1* cells. We found that cells lacking one *TRR1* allele, and cells in which *TRR1* was depleted from *tetO* in doxycycline, consumed oxygen more rapidly than WT cells (Fig. 6C). These cells’ increased respiration, in the setting of decreased upper glycolytic flux, might be fueled by pyruvate that is generated from glyceraldehyde-3-P emerging from the accelerated PPP. While we do not know which regulatory systems induce accelerated respiration in *TRR1*-depleted cells, we note that in consequence, these cells must cope with increased formation of reactive oxygen species ^56^ while one of their major oxidative damage repair systems is failing.

### Gluconeogenesis is reduced in *TRR1*-depleted cells

We wondered whether an increased glucose-6-P demand for provision of the accelerated PPP might induce gluconeogenesis in *TRR1*-depleted cells. We assayed the activity of fructose-1,6-bisphosphatase, which catalyzes an irreversible, gluconeogenesis-specific step that produces fructose-6-P, the gluconeogenic precursor of glucose-6-P. As expected, lactate-grown cells of both genotypes had drastically higher fructose-1,6-bisphosphatase activity than glucose-grown cells (Fig. 6D), since they must produce all glucose-derived metabolic intermediates through gluconeogenesis. However, *TRR1*-depleted cells had significantly lower fructose-1,6-bisphosphatase activity than WT cells on both carbon sources (Fig. 6D), indicating decelerated gluconeogenesis in these cells. As noted, glucose-6-P and fructose-6-P levels were both substantially lower in *TRR1*-depleted cells than WT (SI Appendix, Fig. S3B). Inability to induce compensatory gluconeogenesis may further decrease these cells’ access to glucose-6-P, exacerbating competition between the PPP and UDP-glucose biogenesis for glucose-6-P.

### Glutathione is the major compensatory system in *TRR1*-depleted cells

Metabolomics analysis showed that glutathione levels were >4-fold higher in glucose-grown *TRR1*-depleted than in WT cells (SI Appendix, Fig. S5A). Since fungi use other antioxidant molecules like erythro-ascorbic acid ^57^, ergothionine ^58^ or kynurenine ^59^, we questioned whether loss of both glutathione and thioredoxin function would abolish viability in *C. albicans*, or whether this species has other redox buffers that can suppress defects caused by loss of these two systems. We constructed homozygous *gcs1* null mutants whose only remaining *TRR1* allele is under *tetO* control. During depletion of *TRR1* on 1 μg/ml doxycycline, these cells failed to grow even on YPD agar, which provides enough extrinsic glutathione for the glutathione auxotrophy of *gcs1/gcs1* cells. Partial depletion of *TRR1* on 300 ng/ml doxycycline allowed some growth on YPD (SI Appendix, Fig. S5B). We concluded that *C. albicans* does not possess redox buffer systems that can compensate combined loss of glutathione and thioredoxin functions, and that most likely, it is upregulated glutathione cycling that consumes NADPH from the accelerated PPP during *TRR1* depletion.

### UDP-glucose biosynthesis is downregulated in *TRR1*-depleted cells

In order to induce UDP-glucose biosynthesis (as well as synthesis of trehalose and glycogen), phosphoglucomutase must compete with G-6-PD and with phosphohexoseisomerase for glucose-6-P. We asked whether acceleration of the PPP during *TRR1* depletion decreases UDP-glucose biosynthesis. Trehalose levels were in fact decreased in *TRR1*-depleted cells. We measured the activity of the key enzyme in this process, UDP-glucose pyrophosphorylase (Ugp1) ^60^. Glucose-grown *TRR1*-depleted cells had significantly decreased Ugp1 activity compared with WT (Fig. 6E). In lactate, WT had lower UGP activity than in glucose, with no significant difference to *TRR1*-depleted cells. Taken together with metabolomics results and other key enzymes’ activities, this finding supports the idea that accelerated flux of glucose-6-P into the PPP during *TRR1* depletion decreases UDP-glucose biosynthesis. Reduced availability of substrate for beta-1,3- and beta-1,6-glucan synthases consequently results in decreased glucan cell wall content and ultimately in the cell wall stress hypersensitivity of *TRR1*-depleted cells.

## Discussion

This work found that Trr1 is required for *C. albicans’* growth and its TORC1-mediated anabolic activity (Fig. 1 A,B). Trr1 is also required for *C. albicans’* tolerance of superoxide- and peroxide stress (Fig. 2). Growth defects of *TRR1*-depleted cells increase with increasing ambient temperature. Thioredoxin reductase is also essential in the human fungal pathogens *Cryptococcus neoformans* ^12^ and *Aspergillus fumigatus* ^14^. Comparison of *TRR1* versus *GCS1* depletion, which disable the thioredoxin and glutathione systems respectively, led to cells’ severe growth defects in the former versus moderate impairment in the latter (Figs. 1, 2). Our findings indicate a central role for the thioredoxin system in *C. albicans*, with apparently a secondary role for glutathione.

Only for morphogenetic phenotypes did *GCS1* depletion result in more severe defects, while *TRR1* depletion led to dysregulated hyphal growth responses to ambient pH (Fig. 1C), suggesting distinct morphogenetic client proteins repaired by thioredoxin versus glutathione in *C. albicans*.

Defective Hog1 activation in *TRR1*-depleted cells may be due to a lack of functional thioredoxin Trx1 in the absence of Trr1, as the former is reported to be essential for Hog1 phosphorylation in response to oxidative stress in *C. albicans* ^5,7,61^. Lack of Trr1 hence not only impairs the key biochemical process to reduce oxidized thiols, but also perturbs one of the cell’s signaling systems that respond to resultant oxidative damage.

Combinatorial stress has been studied for host-related stressors like oxidative and nitrosative stress, cationic, osmotic and nutritional stress as reviewed e.g. in ^62^. Anticipatory stress resistance systems provide cross-protection when distinct stresses are sequentially imposed ^33,34^. Cell wall stress may be exerted by macrophages on hyphal cells ^63^, and importantly, cell wall stress is the cidal mechanism of the first-line medications for invasive candidiasis, echinocandins ^64^. Imposition of oxidative stress by depleting *TRR1* was not cross-protective against cell wall stress; *TRR1*-depleted cells were hypersensitive to inhibition of beta-1,3-glucan synthesis and to disruption of chitin fibrils (Fig. 3A). Conversely, cells experiencing genetically induced cell wall stress, in which essential cell wall biosynthetic enzymes Kre6 and Chs1 were depleted genetically, were then hypersensitive to the thiol-oxidative stressor diamide (Fig. 3C), indicating that mutual potentiation of cell wall- and oxidative stress could be a more general phenomenon.

Mkc1 phosphorylation, indicating activation of the cell wall integrity pathway, was defective in response to cell wall stress during *TRR1* transcriptional dysregulation. Navarro-Garcia and colleagues previously found that Mkc1 phosphorylation in response to oxidative and nitrosative-, but not to cell wall stress, depends on Hog1 ^36^. A mechanism by which Trr1 is required for appropriate Mkc1 activation during cell wall stress currently remains enigmatic.

Increased cell wall stress hypersensitivity of *TRR1*-depleted cells on lactate raised the question whether the bulk of structurally stabilizing cell wall polymers ^41^, beta-1,3- and beta-1,6-glucan, are produced normally. In fact, *TRR1* depletion led to decreased cell wall glucan content (Fig. 4B).

Chitin comprises a much smaller fraction of *C. albicans’* cell wall polysaccharides ^41^ but its production is induced during cell wall stress in cells with intact Mkc1 signaling ^48,49^. We found increased cell wall chitin in *TRR1*-depleted cells despite their defective Mkc1 signaling (Fig. 4C). IN-acetylglucosamine, a precursor in chitin biosynthesis, was actually increased in *TRR1*-depleted cells (SI Appendix, Fig. S3B). These findings together suggest as yet unknown cell wall stress-responsive signaling systems in *C. albicans*.

Why do *TRR1*-depleted cells produce less cell wall glucans? Our metabolomics experiments together with our enzymatic assays are consistent with increased flux through the PPP in these cells (Fig. 6A), which in yeast and animal cells is a rapid response to oxidative stress ^50,51,55,65^.

Concomitantly decreased glucose-6-P flux into glycolysis is a general feature of this carbon metabolism reconfiguration, which our assay of phosphofructokinase activity confirmed for *TRR1*-depleted cells (Fig. 6B). But a third pathway needs glucose-6-P: UDP-glucose biosynthesis as a precursor for glycogen, trehalose and, in ascomycetes, cell wall beta-1,3- and beta-1,6-glucans (Fig. 5A). For these fungi, preferential allocation of precious glucose-6-P for NADPH generation to manage oxidative stress while deprioritizing cell wall construction, may be adaptive: oxidative damage to essential proteins and DNA could end in death or replication failure, while cells with structurally weak cell walls might survive. Peroxide stress was found by others to lead to remodeling of carbon metabolism-involved protein and phosphoprotein abundance ^66^.

Unexpectedly, gluconeogenesis was diminished in *TRR1*-depleted cells as evinced by their decreased fructose-1,6-bisphosphatase activity (Fig. 6D). Given increased need for glucose-6-P in these cells for accelerated flux into the PPP, increased gluconeogenesis would appear appropriate. *TRR1*-depleted cells’ failure to induce gluconeogenesis may be mechanistically related to an observation in *S. cerevisiae*: during oxidative stress, the peroxiredoxin Tsa1 binds and downregulates pyruvate kinase Pyk1 which catalyzes the last and irreversible step in glycolysis, saving glycolytic intermediates for use in gluconeogenesis ^67^. *C. albicans* has a Tsa1 homolog, with 73% identity at the amino acid level ^68^, which is a major substrate of *C. albicans* thioredoxin ^5^. We speculate that depletion of *TRR1* suspends Tsa1 in an oxidized state such that it can no longer bind and inhibit Pyk1, resulting in inefficient upregulation of gluconeogenesis. Regardless of the mechanism, inability to adequately upregulate gluconeogenesis likely exacerbates *TRR1*-depleted cells’ metabolic pathway competition for glucose-6-P and further compromises cell wall glucan production.

In summary, in addition to defective direct repair of oxidatively damaged cellular components, lack of Trr1 leads to paradoxical failures of important regulatory systems: the HOG pathway, the cell wall integrity pathway, and gluconeogenesis. Other systems we did not characterize may also be dysregulated, given the importance of redox regulation of signaling systems and enzymatic activities ^69^. As loss of Trr1 activity sensitizes *C. albicans* to agents from the 2 fungicidal drug classes in clinical use, the polyene AmphotericinB and the echinocandin micafungin, pharmacological inhibition of Trr1 could alone or in combination, have potent anticandidal effects. Since fungal thioredoxin reductases are highly conserved but structurally completely different from their human counterparts (SI Appendix, Fig. S1), small molecule inhibitors of Trr1 may have a broad antifungal spectrum and low human toxicity.

## Methods

All experiments were performed for ≥3 biological replicates on different days. Error bars in figures depict standard deviations; statistical significance marks used the following criteria: ns is p>0.05, * 0.01<p≤0.05, ** 0.001<p≤0.01, *** 0.0001<p≤0.001. Detailed methods are included in SI Appendix.

### Strains and culture conditions

The *C. albicans* strains used are described in SI Appendix Table S1. *C. albicans* strains were generated in the SC5314 genetic background using the *CaNAT1* selectable marker and the *tetO* regulatory element in cognate plasmid constructs (SI Appendix Table S2), as described in ^70^. At least two independent heterozygotes were used to derive all mutants, and their identical phenotypes were confirmed before one series was chosen for further characterization. Mutations were confirmed by PCR spanning the upstream and downstream homologous recombination junctions of transforming constructs, and sequencing of PCR products using primers listed in SI Appendix Table S3. Media were used as in ^38^. Cells were freshly recovered for 2 days from frozen stock for each experiment. *Tet-O*-controlled mutants in *TRR1* and *GCS1* were recovered on medium containing the final doxycycline concentration for each experiment except metabolomics, since functional depletion of Trr1 and of reduced thioredoxin apparently occurred over several cell divisions. Incubation was at 30^°^C, except morphogenesis phenotyping at 37^°^ and AmphotericinB Etest at 35^°^.

### Stress phenotypes and growth on distinct carbon sources or amino acids

Recovered cells were washed and diluted with 0.9% NaCl in 3-fold steps starting with OD_600_ 0.5. Cell suspensions were pinned onto agar media by replicator with prongs calibrated to 1.5 μl (V&P Scientific, VP407). For stress phenotypes and amino acid auxotrophy testing, 1 μg/ml doxycycline while for distinct carbon sources, 300 ng/ml doxycycline were used to repress gene expression. Growth was quantified for histogram visualization by measuring spot signal intensity in ImageJ (imagej.net/welcome).

### Western Blots

Cell harvesting, lysis and Western blotting were performed as described in ^17^ using antibodies listed in SI Appendix Table S4. For densitometry, ImageJ was used as in ^17^.

### Hyphal morphogenesis assay

Recovered cells were washed and resuspended in 0.9% NaCl to OD_600_ 0.1. Variations between single colonies and colony density effects were minimized by spotting 3 μl cell suspension at 6 equidistant points, using a template, around the perimeter of an agar plate. RPMI plates were buffered to pH 5 with 100 mM MES and to pH 7 with 165 mM MOPS and incubated 2-4 days at 37°.

### Etest

YPD-recovered cells were washed and resuspended in 0.9% NaCl to OD_600_ 2. A sterile swab (Puritan, 25-806 1WC) was soaked in the cell suspension and after removal of excess fluid, swabbed in three directions on RPMI 1640 agar buffered to pH 7 with 165 mM MOPS. Swabbing was repeated twice; plates were dried, applied with AmphotericinB Etest strips (Liofilchem, 92153) and incubated at 35°C for 24 to 48 hours.

### Growth curves

Cells were washed in 0.9% NaCl and diluted to OD_600_ 0.1 in 150 μl medium in flat bottom 96-well plates. OD_600_ readings were monitered every 15 minutes in a BioTek Synergy 2 Multi-Mode Microplate Reader (Winooski, VT, USA). Standard deviations of 3 technical replicates, representing separate wells, were calculated, and graphed in GraphPad Prism Version 9.1.0 (216), and displayed as error bars.

### Beta-glucan cell wall content measurement

Cells were recovered on SC agar containing 1 μg/ml doxycycline, with either 2% glucose or 2% lactate, grown at 30^°^ C for 1 day, passaged on the same medium and grown for another day. ß-glucan cell wall content was measured from plate-grown cells using β-Glucan Assay Kit (Yeast and Mushroom) (Megazyme by Neogen, K-YBGL) according to the manufacturer’s instructions.

### Chitin cell wall content measurement

Cells used for ß-glucan cell wall content measurement were also processed for chitin measurement as described in ^38^.

### Metabolomics analysis

Recovered WT (JKC915) and *trr1/tetO-TRR1* (JKC2997) cells were grown in liquid SC (300 ng/ml doxycycline) with 2% glucose or 2% lactate for 6 h. Cells were prepared for metabolomics as in ^38^ which was performed as in ^52^. Detailed metabolomics methodology is described in SI Appendix Detailed Methods.

### Enzymatic activity assays

WT (JKC915) and *trr1/tetO-TRR1* (JKC2997) cells were cultured as for metabolomics analysis. Protein concentration of their extracts was measured using BCA assay (Thermo Scientific, 23227). Glucose-6-P dehydrogenase activity was measured using Kit NBP3-24493 (Novus Biologicals). Fructose 1,6-bisphosphatase activity was measured using Kit KTB1331 (Abbkine). Phosphofructokinase activity was measured using Kit NBP3-25908 (Novus Biologicals). Uridine Diphosphate Glucose Pyrophosphorylase activity was measured using Kit E-BC-K857-M (Elabscience); all according to manufacturers’ instructions.

### Oxygen consumption

Cells were grown overnight in SC liquid medium and reinoculated into the same medium; then grown until the logarithmic phase. They were adjusted to 5 x 10^6^ cells/ml in 0.9% NaCl and after addition of glucose oxygen consumption was measured every 5 min. Cultures contained no or 300 ng/ml doxycycline.

## Acknowledgments

This work was funded by R21AI178693 to JRK. We thank Bin Bao for assistance in flow cytometry at the Boston Children’s Hospital Cell Function and Imaging Core. We thank Rebeca Alonso Monge and Jesús Pla, both of Universidad Complutense de Madrid, Spain, and Alessandra Da Silva Dantas, Newcastle University, UK, for helpful discussions.

## Supporting Information

Appendix 1 (PDF):

Supplementary Figures and Figure legends Fig. S1-S5

Detailed Methods

SI Table 1. Strains

SI Table 2. Plasmids

SI Table 3. Oligonucleotides

SI Table 4. Antibodies

Dataset S1 Metabolomics Results

