## Supplemental Information for "Metabolic pathway competition sensitizes thioredoxin reductase-depleted *Candida albicans* to cell wall stress and antifungals"

### Supplementary Figures

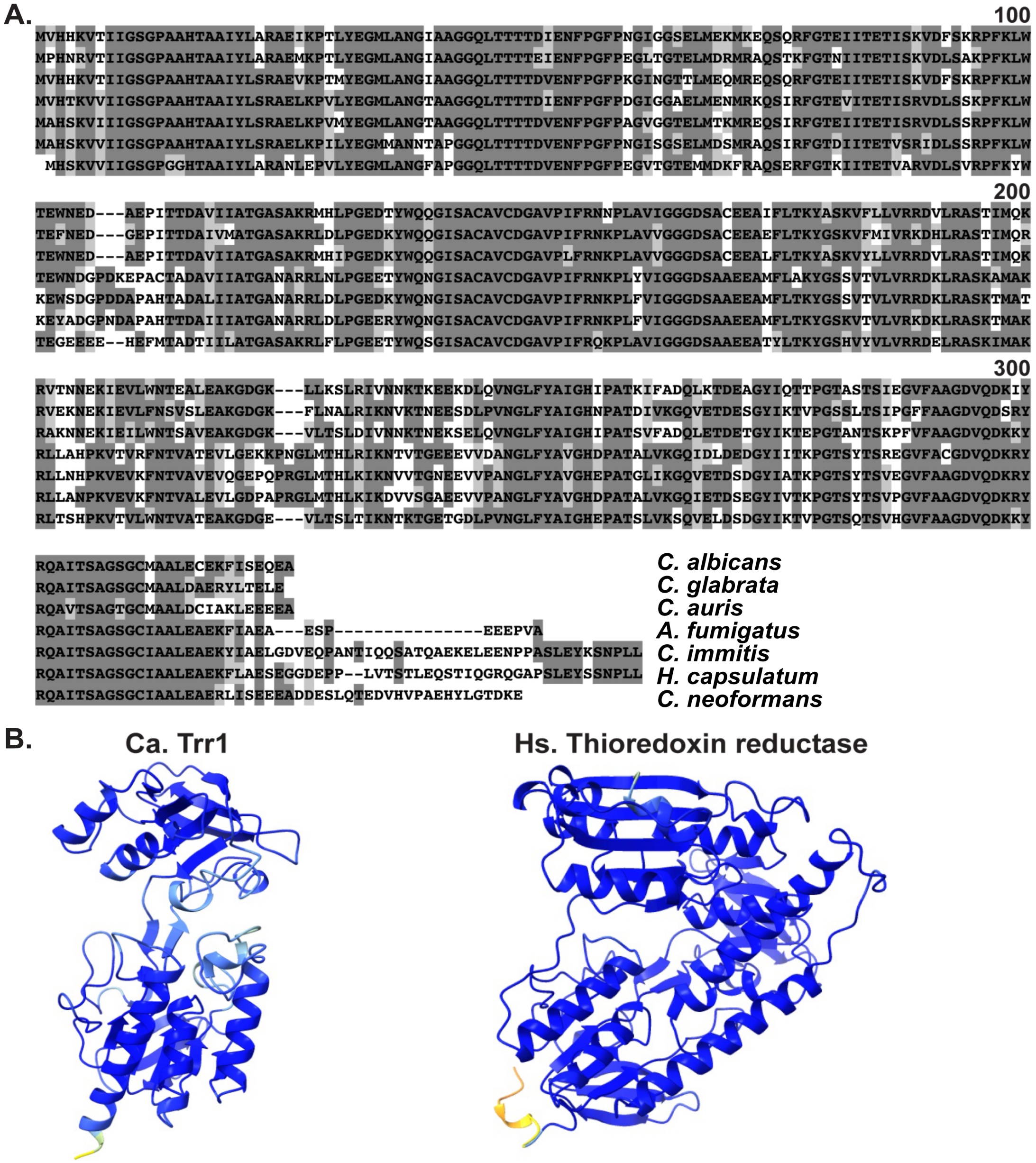


**Figure S1. Thioredoxin reductases of human-pathogenic fungi are conserved, while the human enzyme is structurally divergent. A.** Primary amino acid sequences of *Candida albicans* thioredoxin reductase and thioredoxin reductase homologs of other important human fungal pathogens. **B.** Alpha-fold protein structure comparison of *Candida albicans* thioredoxin reductase (Ca. Trr1) and *Homo sapiens* (Hs.) Thioredoxin reductase.

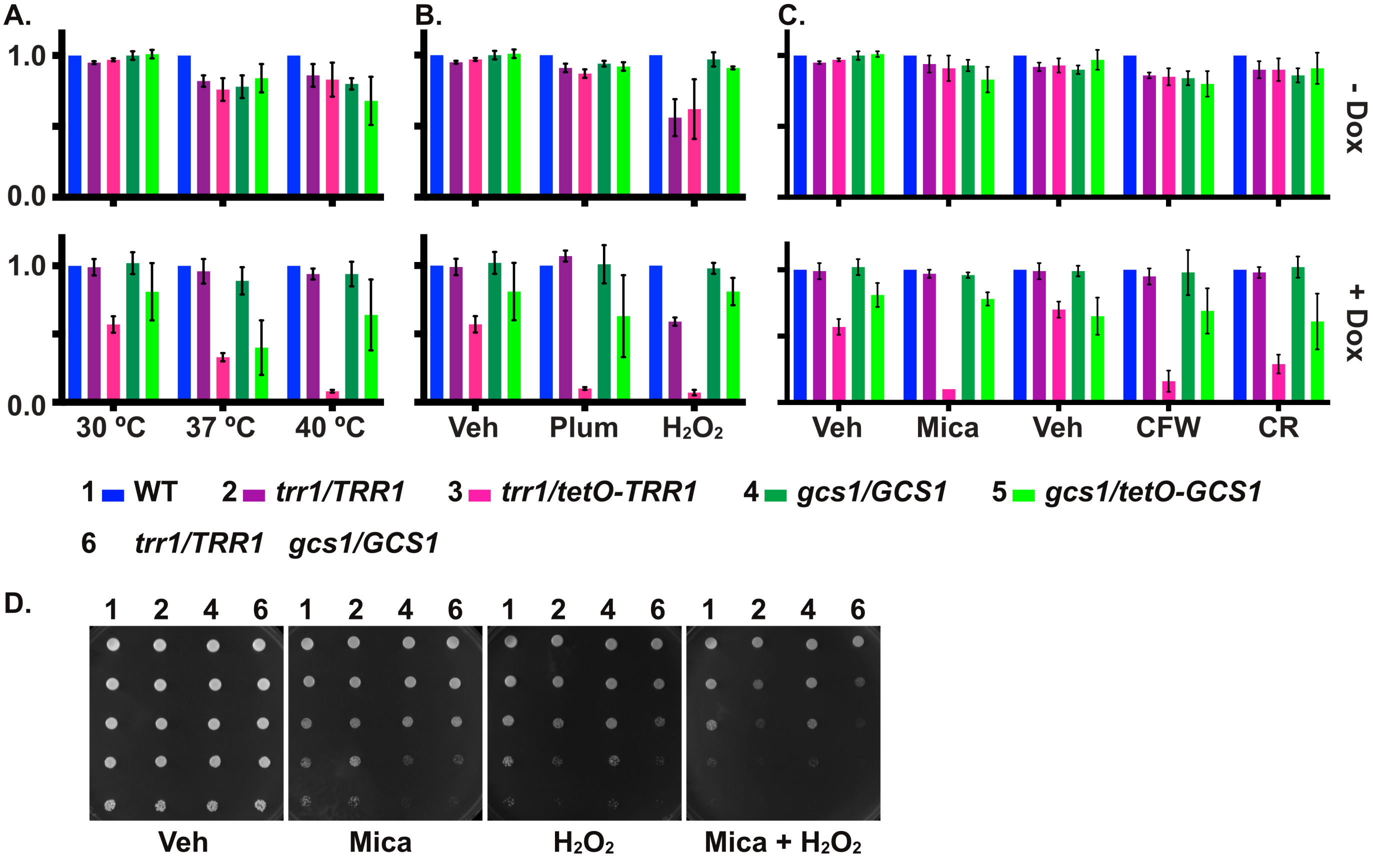


**Figure S2. Loss of Trr1 activity sensitizes cells to oxidative and cell wall stress, and these stresses potentiate each other even in hemizygous cells. A-C.** Growth quantitation of *TRR1* and *GCS1* depleted cells: histograms of Fig. 1A, Fig. 2A, Fig. 3A cell suspension dilution spots, respectively. Growth calculated in ImageJ based on imaged spots' intensity and normalized to wild type (WT) growth in each condition. Error bars: SD of 3 biological replicates. **D.** Heterozygous phenotypes comparing wild type with *trr1/TRR1, gcs1/GCS1* and *trr1/TRR1 gcs1/GCS1* cells: dilutions (top to bottom) of indicated genotypes spotted onto SC plates containing vehicle (Veh, $H_2O$) or 15 ng/ml micafungin (Mica), 2 mM $H_2O_2$ or combined stresses, grown at 30$^o$ C for 1 day. Strains are 1. WT (JKC915); 2. *trr1/TRR1* (JKC2989); 3. *trr1/tetO-TRR1* (JKC2997); 4. *gcs1/GCS1* (JKC3001); 5. *gcs1/tetO-GCS1* (JKC3009). 6. *trr1/TRR1 gcs1/GCS1* (JKC3508).

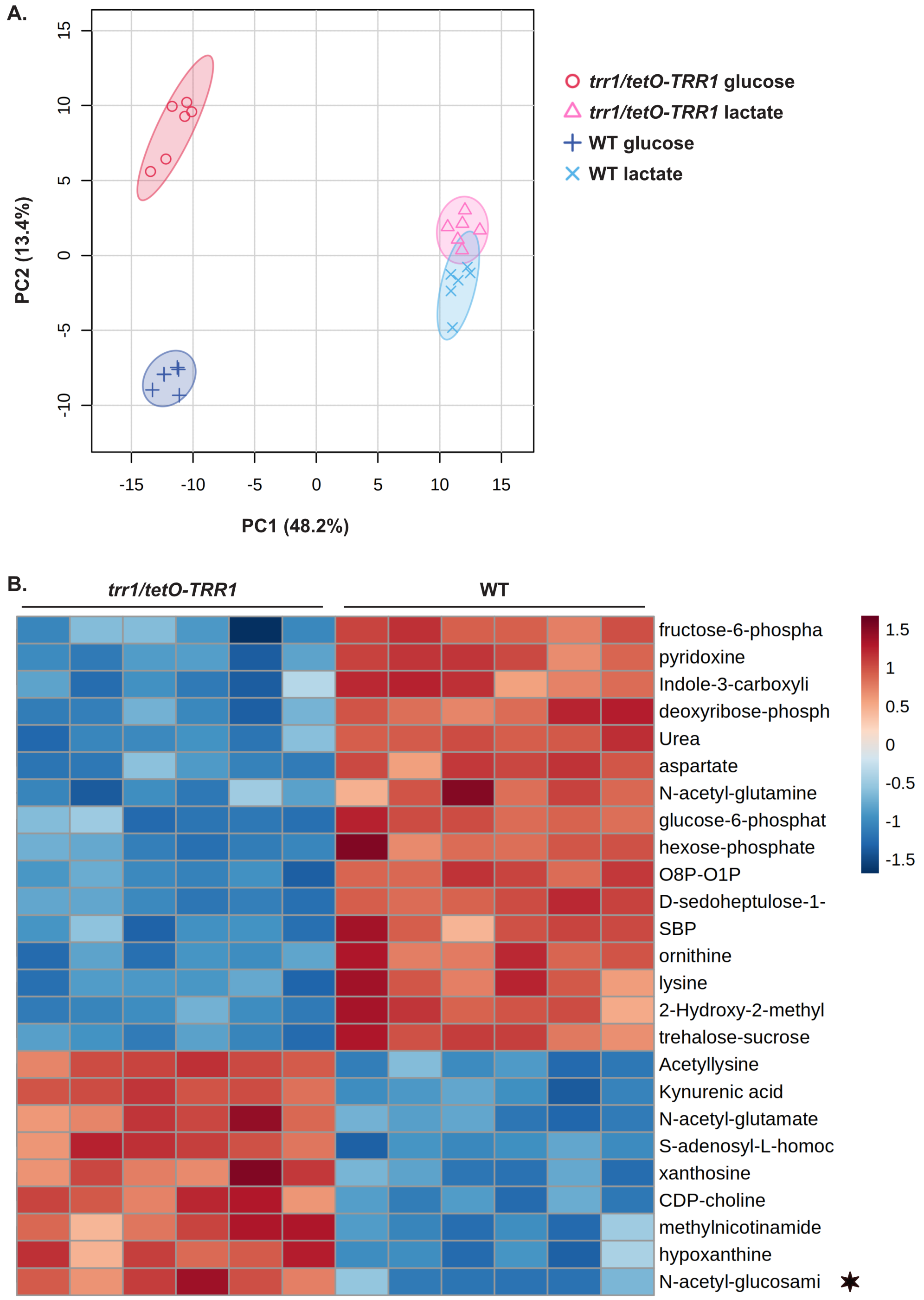


**Figure S3. Metabolomic profiles of glucose-grown *TRR1*-depleted versus WT cells are distinct. A.** PCA of metabolomic profiles from wild type (WT) and *trr1/tetO-TRR1* cells grown in glucose or lactate with 300 ng/ml doxycycline. PCA shows clear separation between *TRR1*-depleted and wild-type cells in glucose-grown samples, while lactate-grown samples displayed closer clustering. **B.** Heatmap of the top 25 metabolites differing most significantly between *trr1/tetO-TRR1* and WT cells grown in glucose, ranked by p-values from unpaired parametric t-tests. *N-acetylglucosamine. Strains are WT (JKC915); *trr1/tetO-TRR1* (JKC2997).

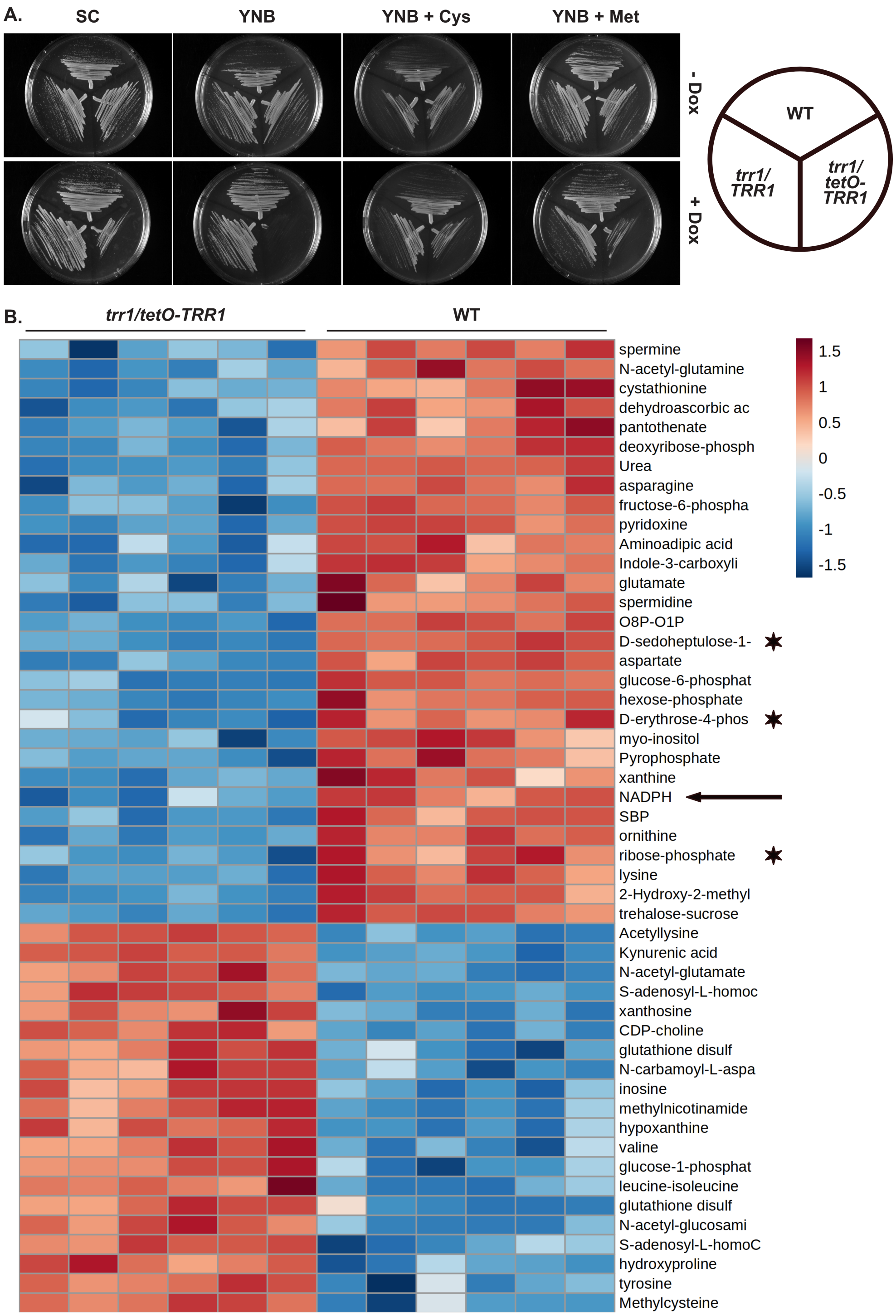


**Figure S4. A. *TRR1*-depleted cells are auxotrophic for cysteine and methionine, and PPP intermediates are among the Top50 significantly different metabolites between glucose-grown *TRR1*-depleted and WT cells. A.** Cells of indicated genotypes were streaked onto SC, YNB, YNB supplemented with 0.125 mg/ml cysteine (Cys) or methionine (Met) plates, without or with 1 μg/ml doxycycline (Dox), grown at $30^{\circ}$ C for 1 day. **B.** Heatmap of the top 50 metabolites differing most significantly between *trr1/tetO-TRR1* and WT cells grown in glucose, ranked by p-values from unpaired parametric t-tests. Arrow: NADPH; asterisks: PPP intermediates. Strains are WT (JKC915); *trr1/TRR1* (JKC2989); *trr1/tetO-TRR1* (JKC2997).

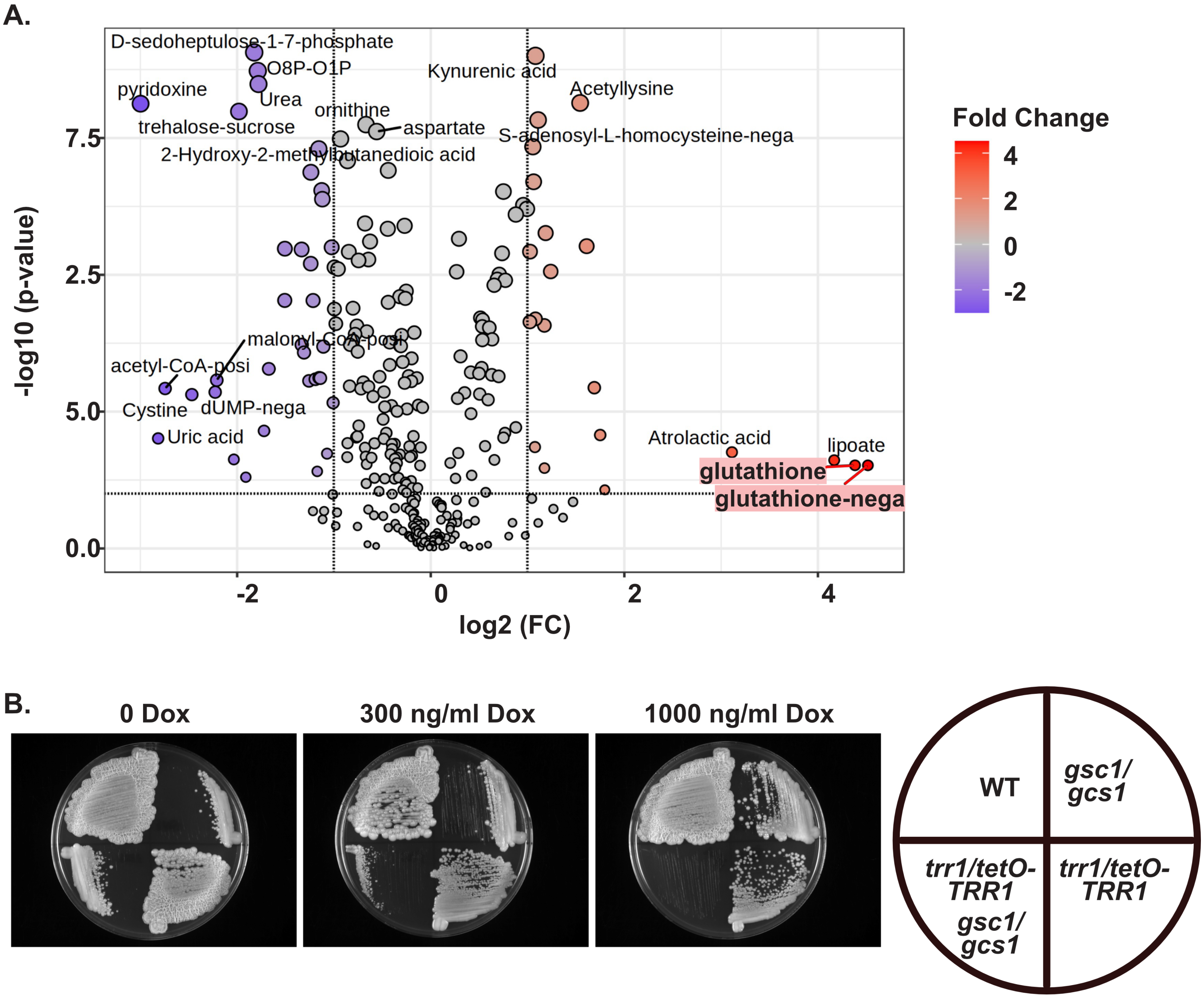


**Figure S5. Glutathione levels are upregulated in *TRR1*-depleted cells, while cells lacking glutathione and maximally repressed for *TRR1* expression, cease growth. A.** Volcano plot of metabolite abundance differences between *TRR1*-depleted and WT cells grown in glucose with 300 ng/ml doxycycline. The x-axis represents $\log_2$ fold change, and the y-axis represents $-\log_{10}$ *p*-value from unpaired t-tests. **B.** Cells of indicated genotypes streaked onto YPD containing doxycycline (Dox) at indicated concentrations, grown at 30° C for 2 days and room temperature for additional 4 days. Strains are WT (JKC915); *gcs1/gcs1* (JKC3491); *trr1/tetO-TRR1* (JKC2997); *gcs1/gcs1 trr1/tetO-TRR1* (JKC3501).

### Detailed Methods

**Stress phenotypes and growth on distinct carbon sources**. Cells recovered from glycerol stock were grown on YPD agar with and without 1 µg/ml (stress phenotypes) or 300 ng/ml (alternative carbon sources) doxycycline at 30° for 36 to 48 hours, then cells were washed in 0.9% NaCl and diluted with 0.9% NaCl in 3-fold steps (unless otherwise stated) from a starting $OD_{600}$ of 0.5 in a microtiter plate. The cell suspensions were pinned onto appropriate agar media using a replicator with prongs calibrated to deliver 1.5 µl (V&P Scientific, VP407). Plates were imaged after 2-3 days of incubation at 30° unless stated otherwise. All panels shown represent at least 3 biological replicates. Growth of each strain was calculated in ImageJ based on imaged spots intensity, normalized to WT growth in each condition and plotted as histograms; error bars show standard deviation (SD) of 3 biological replicates.

**Western Blots.** Cells were revived from frozen stocks on solid YPD agar without or with 1 µg/ml doxycycline for 2 days. Cells were washed in 0.9% NaCl and inoculated into 200 ml YPD without or with 300 ng/ml doxycycline to an $OD_{600}$ of 0.3 and pre-grown at 30°, 200 rpm for 3.5 hours. Cells from pre-growth were collected at 2000 rpm for 5 minutes, washed in 0.9% NaCl and re-inoculated at $OD_{600}$ of 0.4 into 50 ml YPD or YPD + 300 ng/ml doxycycline with vehicle ($H_2O$) or 10 ng/ml micafungin or with 2.5 mM $H_2O_2$, and grown at 30° (30 minutes for vehicle and micafungin, 5 minutes for $H_2O_2$). Treated cells were collected, washed twice with ice cold water and resuspended in lysis buffer. Cells were lysed with silicon beads using a beads beater. Total protein concentration was quantified using the Pierce™ BSA Protein Assay Kit (Thermo Scientific, 23225). 25 µg of total protein was separated by SDS-PAGE and transferred to PVDF membranes, then probed for phospho-S6 using anti-phospho (S/T) Akt substrate rabbit polyclonal antibody (Cell Signaling Technology, 9611), phospho-Hog1 using anti-Phospho-p38 MAPK (Thr180/Tyr182) (D3F9) rabbit monoclonal antibody (Cell Signaling Technology, 4511), phospho-Mkc1 using anti-Phospho-p44/42 MAPK (Erk1/2) (Thr202/Tyr204) rabbit monoclonal antibody (Cell Signaling Technology, 4370), and anti-PSTAIR mouse monoclonal antibody (Sigma Aldrich Inc, SAB4200861) as loading control. Secondary antibodies used were anti-rabbit IgG, HRP-linked antibody (Cell Signaling Technology, 7074) and anti-mouse IgG, HRP-linked antibody (Cell Signaling Technology, 7076). The images were developed using SuperSignal™ West Pico Plus Chemiluminescent Substrate (Thermo Scientific, 34580) and were imaged using Azure Biosystems C600 gel Imaging System. Densitometry was calculated using ImageJ (imagej.net/welcome) analysis software (opensource).

**Etest.** Cells recovered from glycerol stock were grown on YPD agar without and with 1 µg/ml doxycycline at 30˚° for 36 to 48 hours, washed and resuspended in 0.9% NaCl to $OD_{600}$ of 2. A non-toxic, sterile swab (Puritan, 25-806 1WC) was soaked in the cell suspension and pressed against the inner wall of the test tube to remove excess fluid. The cells were swabbed on RPMI 1640 media buffered to pH 7 with 165 mM MOPS, covering three directions to evenly distribute the inoculum. The swabbing was repeated twice. The plates were allowed to completely dry and then AmphotericinB (AMB 0.002-32 mg/L) Etest strips (Liofilchem, 92153) were placed. The plates were incubated at 35° for 24 to 48 hours.

**Chitin measurement.** Cells from the cultures used for beta-glucan cell wall content measurement were fixed with formaldehyde (Sigma, F-8775, final fixing concentration at 1.8%) at room temperature for 30 min and washed twice with Phosphate Buffered Saline (pH 7.4), then stained with 0.01 mg/mL Calcofluor White (Sigma, F-3543-1G) prepared in $H_2O$ for 4 min at room temperature. Stained cells were washed five times with 0.9% NaCl and resuspended in 500 µL 0.9% NaCl. Calcofluor White stained cells were vortexed before processing with flow-cytometer (BD FACS LSRFortessa, BD Biosciences, 647796L6). Chitin staining was detected using the Pacific Blue channel with violet Laser (405 nm) and signal collected with channel Pacific Blue (fluorescence filter 450/40 nm). $10^5$ events were recorded for data analysis.

**Metabolomics analysis.** WT (JKC915) and *trr1/tetO-TRR1* (JKC2997) cells were recovered on SC plates containing 1 µg/ml doxycycline, with either 2% glucose or 2% lactate as carbon source and grown at 30° for 1 day. Cells were washed in 0.9% NaCl and inoculated into SC medium containing 300 ng/ml doxycycline, with either 2% glucose or 2% lactate as carbon source, with final $OD_{600}$ at 0.2 and grown at 30°, 200 rpm for 6 h. Harvested cells were washed with ice-cold water 3 times and cell pellets were resuspended in 500 µl 80% methanol (cooled at -80°) on ice and transferred to -80° for 40 min. Metabolites were extracted by centrifugation at 4°, top speed for 5 min. Pellets were re-extracted twice by resuspension in 400 µl 80% methanol and centrifugation. All supernatants were combined and dried completely in an Eppendorf 5301 Vacufuge Concentrator (Eppendorf, USA). The dried samples were used for metabolomics analysis as in [1].

Samples were re-suspended using 20 µl HPLC grade water for mass spectrometry. 5-7 µL were injected and analyzed using a hybrid 6500 QTRAP triple quadrupole mass spectrometer (AB/SCIEX) coupled to a Prominence UFLC HPLC system (Shimadzu) via selected reaction monitoring (SRM) of a total of 300 endogenous water soluble metabolites for steady-state analyses of samples. Some metabolites were targeted in both positive and negative ion mode for a total of 311 SRM transitions using positive/negative ion polarity switching. ESI voltage was +4950V in positive ion mode and –4500V in negative ion mode. The dwell time was 3 ms per SRM transition and the total cycle time was 1.55 seconds. Approximately 9-12 data points were acquired per detected metabolite. Samples were delivered to the mass spectrometer via hydrophilic interaction chromatography (HILIC) using a 4.6 mm i.d x 10 cm Amide XBridge column (Waters) at 400 µl/min. Gradients were run starting from 85% buffer B (HPLC grade acetonitrile) to 42% B from 0-5 minutes; 42% B to 0% B from 5-16 minutes; 0% B was held from 16-24 minutes; 0% B to 85% B from 24-25 minutes; 85% B was held for 7 minutes to re-equilibrate the column. Buffer A was comprised of 20 mM ammonium hydroxide/20 mM ammonium acetate (pH=9.0) in 95:5 water:acetonitrile. Peak areas from the total ion current for each metabolite SRM transition were integrated using MultiQuant v3.0.2 software (AB/SCIEX).

For data analysis, MetaboAnalystR package (v 4.0) was used to preprocess raw metabolite quantitation data and perform pairwise comparison analysis between different experimental conditions [2]. Each dataset was processed with sum normalization, log transformation, and auto-scaling, and missing values were imputed using minimum value replacement. Statistical analyses included unpaired parametric t-tests and fold change analysis, and the metabolites were ranked by the t-test p values to identify those most differentiating *TRR1*-depleted and WT cells. Differentially abundant metabolites were identified using an absolute log2-fold change ≥ 1 and an unpaired parametric t-test with FDR < 0.05, and were further used for pathway enrichment analysis in MetaboAnalyst (v 6.0) web platform using the KEGG database (species: *C. albicans*), and the pathway impact scores were calculated to integrate both the number of significantly altered metabolites and their topological importance within each pathway, assigning higher scores to changes occurring at highly connected or central nodes [3]. Barplots show enriched KEGG pathways for multiple pairwise comparisons, displaying pathways with $p < 0.05$ and pathway impact score > 0.

**Enzymatic activities.** WT (JKC915) and *trr1/tetO-TRR1* (JKC2997) cells were cultured as for metabolomics analysis. Proteins were extracted from harvested cells and protein concentration was measured using BCA assay (Thermo Scientific, 23227). Glucose-6-P dehydrogenase activity was measured using Glucose 6 Phosphate Dehydrogenase Activity Assay Kit (Colorimetric) (Novus Biologicals, NBP3-24493). Fructose 1,6-bisphosphatase activity was measured using CheKine™ Micro Fructose 1,6-Bisphosphatase (FBP) Activity Assay Kit (Abbkine, KTB1331). Phosphofructokinase activity was measured using Phosphofructokinase/PFK Activity Assay Kit (Colorimetric) (Novus Biologicals, NBP3-25908). Uridine Diphosphate Glucose Pyrophosphorylase activity was measured using Elabscience® Uridine

Diphosphate Glucose Pyrophosphorylase (UGP) Activity Assay Kit (Elabscience, E-BC-K857-M). All assays were performed according to manufacturers' instructions.

**Oxygen consumption.** Cells were recovered on YPD agar containing 1 µg/ml doxycycline and grown at 30° for 1 day. Isolated colonies of each strain were inoculated into SC liquid medium and grown overnight at 30° (200 rpm) without or with 300 ng/ml doxycycline. Aliquots of the overnight culture were inoculated into fresh SC medium without or with 300 ng/ml doxycycline and incubated at 30° (200 rpm) until the logarithmic growth phase was reached (approximately 3 h). Cells from logarithmic growth phase were washed with 0.9% NaCl without or with 300 ng/ml doxycycline and adjusted to approximately 5 x $10^6$ cells/ml. Oxygen consumption was measured every 5 min in a 700 µl suspension prepared in 0.9% NaCl without or with 300 ng/ml doxycycline, measurement started with the addition of glucose (to achieve a 2% final glucose concentration). Oxygen consumption was measured using a Clark-type electrode (dual digital-model 20; Rank Brothers. Ltd., Cambridge, United Kingdom) at 25°, according to the manufacturer's instructions.

**SI Table 1.** Strains used in this study

| ***C. albicans* strain name** | **Parent** | **Genotype** | **Strain construction** | **Reference** |
|---|---|---|---|---|
| SC5314 | | Wild type | | [4] |
| JKC915 | SC5314 | *HIS1/his1:: tetR-FRT* | | [5] |
| JKC2977 | JKC915 | *HIS1/his1:: tetR-FRT TRR1/trr1::uPAM-FRT-FLP-NAT1* | JKC915 transformed with KpnI/BsiWI digested pJK1554 to knockout the 1st allele of *TRR1*. | This work |
| JKC2989 | JKC2977 | *HIS1/his1:: tetR-FRT TRR1/trr1::uPAM-FRT* | JKC2977 *NAT1* flipped out | This work |
| JKC2936 | JKC915 | *HIS1/his1:: tetR-FRT TRR1/trr1::FRT-FLP-NAT1-tetO-TRR1* | JKC915 transformed with KpnI/NcoI digested pJK1557 to have one of the *TRR1* alleles under *tetO* control. | This work |
| JKC2942 | JKC2936 | *HIS1/his1:: tetR-FRT TRR1/trr1::FRT-tetO-TRR1* | JKC2936 *NAT1* flipped out | This work |
| JKC2981 | JKC2942 | *HIS1/his1:: tetR-FRT trr1::uPAM-FRT-FLP-NAT1/trr1::FRT-tetO-TRR1* | JKC2942 transformed with KpnI/BsiWI digested pJK1554 to knock out the WT allele of *TRR1* and have the only allele of *TRR1* under *tetO* control. | This work |
| JKC2997 | JKC2981 | *HIS1/his1:: tetR-FRT trr1::uPAM-FRT /trr1::FRT-tetO-TRR1* | JKC2981 *NAT1* flipped out | This work |
| JKC2982 | JKC2942 | *HIS1/his1:: tetR-FRT trr1::uPAM-FRT-FLP-NAT1/trr1::FRT-tetO-TRR1* | JKC2942 transformed with KpnI/BsiWI digested pJK1554 to have the only allele of *TRR1* under *tetO* control. | This work |
| JKC2999 | JKC2982 | *HIS1/his1:: tetR-FRT trr1::uPAM-FRT /trr1::FRT-tetO-TRR1* | JKC2982 *NAT1* flipped out | This work |
| JKC2983 | JKC915 | *HIS1/his1:: tetR-FRT GCS1/gcs1::uPAM-FRT-FLP-NAT1* | JKC915 transformed with KpnI/BsiWI digested pJK1555 to knockout the 1st allele of *GCS1*. | This work |
| JKC3001 | JKC2983 | *HIS1/his1:: tetR-FRT GCS1/gcs1::uPAM-FRT* | JKC2983 *NAT1* flipped out | This work |
| JKC2939 | JKC915 | *HIS1/his1:: tetR-FRT GCS1/gcs1::FRT-FLP-NAT1-tetO-GCS1* | JKC915 transformed with KpnI/NcoI digested pJK1559 to have one of the *GCS1* alleles under *tetO* control. | This work |
| JKC2948 | JKC2939 | *HIS1/his1:: tetR-FRT GCS1/gcs1::FRT-tetO-GCS1* | JKC2939 *NAT1* flipped out | This work |
| JKC2987 | JKC2948 | *HIS1/his1:: tetR-FRT gcs1::uPAM-FRT-FLP-NAT1/gcs1::FRT-tetO-GCS1* | JKC2948 transformed with KpnI/BsiWI digested pJK1555 to knock out the WT allele of *GCS1* and have the only allele of *GCS1* under *tetO* control. | This work |
| JKC3009 | JKC2987 | *HIS1/his1:: tetR-FRT gcs1::uPAM-FRT /gcs1::FRT-tetO-GCS1* | JKC2987 *NAT1* flipped out | This work |
| JKC3456 | JKC915 | *HIS1/his1:: tetR-FRT gcs1::uPAM-FRT-FLP-NAT1-FRT/GCS1* | JKC915 transformed with KpnI/BsiWI digested pJK1638 to knock out the 1st allele of *GCS1*. | This work |
| JKC3463 | JKC3456 | *HIS1/his1:: tetR-FRT gcs1::uPAM-FRT/GCS1* | JKC3456 *NAT1* flipped out | This work |
| JKC3475 | JKC3463 | *HIS1/his1:: tetR-FRT gcs1::uPAM-FRT/gcs1::uPAM-FRT-FLP-NAT1-* | JKC3463 transformed with KpnI/BsiWI digested pJK1555 to knock out the 2nd allele | This work |

| | | | | |
|---|---|---|---|---|
| | | *FRT* | of *GCS1*. | |
| JKC3491 | JKC3475 | *HIS1/his1:: tetR-FRT gcs1::uPAM-FRT/gcs1::uPAM-FRT* | JKC3475 *NAT1* flipped out | This work |
| JKC3461 | JKC2997 | *HIS1/his1:: tetR-FRT trr1::uPAM-FRT/trr1::FRT-tetO-TRR1 gcs1::uPAM-FRT-FLP-NAT1-FRT/GCS1* | JKC2997 transformed with KpnI/BsiWI digested pJK1638 to knockout the 1st allele of *GCS1* in the *trr1/tetO-TRR1* background. | This work |
| JKC3473 | JKC3461 | *HIS1/his1:: tetR-FRT trr1::uPAM-FRT/trr1::FRT-tetO-TRR1 gcs1::uPAM-FRT/GCS1* | JKC3461 *NAT1* flipped out | This work |
| JKC3484 | JKC3473 | *HIS1/his1:: tetR-FRT gcs1::uPAM-FRT/gcs1::uPAM-FRT-FLP-NAT1-FRT trr1::uPAM-FRT/trr1::FRT-tetO-TRR1* | JKC3473 transformed with KpnI/BsiWI digested pJK1555 to knock out the 2nd allele of *GCS1*, making homozygous *gcs1/gcs1* in the *trr1/tetO-TRR1* background. | This work |
| JKC3501 | JKC3484 | *HIS1/his1:: tetR-FRT gcs1::uPAM-FRT/gcs1::uPAM-FRT trr1::uPAM-FRT/trr1::FRT-tetO-TRR1* | JKC3484 *NAT1* flipped out | This work |
| JKC3506 | JKC2989 | *HIS1/his1:: tetR-FRT gcs1::uPAM-FRT-FLP-NAT1-FRT/GCS1 trr1::uPAM-FRT/TRR1* | JKC2989 transformed with KpnI/BsiWI digested pJK1555 to knock out one allele of *GCS1* in the *trr1/TRR1* heterozygote background, making *trr1/TRR1* and *gcs1/GCS1* double heterozygotes. | This work |
| JKC3508 | JKC3506 | *HIS1/his1:: tetR-FRT gcs1::uPAM-FRT/GCS1 trr1::uPAM-FRT/TRR1* | JKC3506 *NAT1* flipped out | This work |

### SI Table 2. Plasmids used in this study.

| Plasmid | Description | Reference |
|---|---|---|
| pJK1000 | *FLP-NAT1 tetO-PES1* construct, vector backbone is pLitmus28 (New England Biolabs) | 5 |
| pJK1372 | *FLP-NAT1 pho87* deletion construct, derived from pJK1364. Product of fjk1846 and r1862 using SC5314 genomic DNA as template was ligated into pJK1364 using KpnI/ApaI sites. | 6 |
| pJK1554 | *FLP-NAT1 trr1* deletion construct, derived from pJK1372. Product of fjk2211 and rjk2212 using SC5314 genomic DNA as template and product of fjk2213 and rjk2214 using SC5314 genomic DNA as template were ligated into pJK1372 using KpnI/AscI and NotI/BsiWI sites, respectively. | This work |
| pJK1555 | *FLP-NAT1 gcs1* deletion construct, derived from pJK1372. Product of fjk2223 and rjk2224 using SC5314 genomic DNA as template and product of fjk2225 and rjk2226 using SC5314 genomic DNA as template were ligated into pJK1372 using KpnI/AscI and NotI/BsiWI sites, respectively. | This work |
| pJK1557 | *FLP-NAT1 tetO-TRR1* construct, derived from pJK1000. Product of fjk2217 and rjk2218 using SC5314 genomic DNA as template and product of fjk2219 and rjk2220 using SC5314 genomic DNA as template were ligated into pJK1000 using KpnI/ApaI and SacII/NcoI sites, respectively. | This work |
| pJK1559 | *FLP-NAT1 tetO-GCS1* construct, derived from pJK1000. Product of fjk2229 and rjk2230 using SC5314 genomic DNA as template and product of fjk2231 and rjk2232 using SC5314 genomic DNA as template were ligated into pJK1000 using KpnI/ApaI and SacII/NcoI sites, respectively. | This work |
| pJK1638 | *FLP-NAT1 gcs1* deletion construct, derived from pJK1372. Product of fjk2229 and rjk2410 using SC5314 genomic DNA as template and product of fjk2225 and rjk2226 using SC5314 genomic DNA as template were ligated into pJK1372 using KpnI/AscI and NotI/BsiWI sites, respectively. | This work |

### SI Table 3. Oligonucleotides used in this study.

| Primer name | Purpose | Sequence 5' to 3' (lower cases - restriction enzyme recognition sites) |
|---|---|---|
| fjk2211 | Forward primer to amplify the *trr1* deletion construct upstream homologous sequence | CATCAAggtaccAAATAAACGAGGGGCCAAGT |
| rjk2212 | Reverse primer to amplify the *trr1* deletion construct upstream homologous sequence | GATggcgcgccGTGACTTTGTGGTGTACCATTGT |
| fjk2213 | Forward primer to amplify the *trr1* deletion construct downstream homologous sequence | TTGGTAAGCAgcggccgcGAACAAGAAGCTTAGATTTTTC AAGAG |
| rjk2214 | Reverse primer to amplify the *trr1* deletion construct downstream homologous sequence | CTCATGcgtacgTGGGGATAGTTCACACCAAA |
| fjk2215 | Forward primer to verify the 5'end of *trr1* deletion mutant | GCCAATGGCGTGATTAGTTT |

| | | |
|---|---|---|
| rjk2216 | Reverse primer to verify the 3'end of *trr1* deletion mutant | ACGTCCTCGTGTTCTCGTTC |
| rjk1339 | Reverse primer to verify the 5'end of *integration of 'FLP-NAT1'* cassette containing constructs | TGGTGTGTTGTTGACAGGCAAC |
| fjk490 | Forward primer to verify the 3'end integration of '*FLP-NAT1'* cassette containing constructs | TCAAGGAGGGTATTCTGGGC |
| fjk1835 | Forward primer to verify the 3'end integration of '*FLP-NAT1-tetO'* constructs | TGTCGTTTCTGATGGGCTTT |
| fjk2217 | Forward primer to amplify the *tetO-TRR1* construct upstream homologous sequence | CATGTGggtaccGGTGGTAGTGCCTGCAAAAA |
| rjk2218 | Reverse primer to amplify the *tetO-TRR1* construct upstream homologous sequence | GATCgggcccCACACACACACACACAAGCCTA |
| fjk2219 | Forward primer to amplify the *tetO-TRR1* construct downstream homologous sequence | GGATCCccgcggATGGTACACCACAAAGTCACTA |
| rjk2220 | Reverse primer to amplify the *tetO-TRR1* construct downstream homologous sequence | CTCATGccatggTGTCTTCCCCTGGTAAATGC |
| fjk2221 | Forward primer to verify the 5'end of *tetO-TRR1* integration | AGCAGCGAGATTCGTCAACT |
| rjk2222 | Reverse primer to verify the 3'end of *tetO-TRR1* integration | CCACCACCAATCACAGCTAA |
| fjk2223 | Forward primer to amplify the *gcs1* deletion construct upstream homologous sequence | CACTCGggtaccTTGTCCTTTTAATCCTTATCTTGGA |
| rjk2224 | Reverse primer to amplify the *gcs1* deletion construct upstream homologous sequence | TATggcgcgccCCCATGATAATACAGTGGTTCCTC |
| fjk2225 | Forward primer to amplify the *gcs1* deletion construct downstream homologous sequence | CTGGTAAGTTgcggccgcGGGATCAAAATAAACAATTAAT AATTAATCGG |
| rjk2226 | Reverse primer to amplify the *gcs1* deletion construct downstream homologous sequence | CTCATAcgtacgCCAGCACCACCAATTAAAGG |
| fjk2227 | Forward primer to verify the 5'end of *gcs1* deletion mutant | GAGACGGAAAGAGGGAGAGG |
| rjk2228 | Reverse primer to verify the 3'end of *gcs1* deletion mutant | CCTGCTCCATCCAACTCACT |
| fjk2229 | Forward primer to amplify the *tetO-GCS1* construct upstream homologous sequence | CATGTAggtaccAGGGATAGGGAGAGGCTCAA |
| rjk2230 | Reverse primer to amplify the *tetO-GCS1* construct upstream homologous sequence | GATAgggcccCCGTCTCGTACCGGAATAAA |
| fjk2231 | Forward primer to amplify the *tetO-GCS1* construct downstream homologous sequence | GGATCCccgcggATGGGTCTTTTATCTATTGGTACACC |
| rjk2232 | Reverse primer to amplify the *tetO-GCS1* construct downstream homologous sequence | CTCATGccatggCGGGGTTGCTTCTACCATAA |
| fjk2233 | Forward primer to verify the 5'end of *tetO-GCS1* integration | GATGCGATTTCATGCGTTTT |
| rjk2234 | Reverse primer to verify the 3'end of *tetO-GCS1* integration | AAACCCTGGTTTGGAAGCAC |
| rjk2410 | Reverse primer to amplify the *gcs1* deletion construct upstream homologous sequence | ATTggcgcgccCCGTCTCGTACCGGAATAAA |

### SI Table 4. Antibodies used in this study.

| **Purpose** | **Antigen recognized** | **Species** | **Source or Reference** |
|---|---|---|---|
| primary | P-S6 | rabbit | Cell Signaling Technology, #9611 |
| primary | P-Mkc1 | rabbit | Cell Signaling Technology, #4370 |
| primary | P-Hog1 | rabbit | Cell Signaling Technology, #4511 |
| loading control | PSTAIRE (Cdc2) | mouse | Millipore Sigma, #SAB4200861 |
| secondary | Rabbit IgG | goat | Cell Signaling Technology, #7074S |
| secondary | Mouse IgG | horse | Cell Signaling Technology, #7076S |
